# Spatial single-cell proteomics defines multicellular niches in the primary prostate cancer microenvironment

**DOI:** 10.64898/2026.04.30.721907

**Authors:** Adriano Martinelli, Andrea Brunello, Melissa Ensmenger, Sofia Karkampouna, Francesco Bonollo, Eva Comperat, Andrea Lunardi, Beat Roth, Martin Spahn, George N. Thalmann, Elena Brunner, Elisabeth Damisch, Lukas Nommensen, Natalie Sampson, Marianna Rapsomaniki, Marianna Kruithof-de Julio

## Abstract

Prostate cancer displays substantial clinical and histopathological heterogeneity which is not fully captured by conventional Gleason grading. To resolve the spatial and phenotypic complexity of the prostate tumor microenvironment, we performed imaging mass cytometry using a prostate-tailored 34-plex antibody panel on a clinically annotated tissue microarray cohort of 195 patients of primary stage disease after radical prostatectomy (523 regions of interest; 2.19 million cells). We identified 34 distinct cell types spanning epithelial, endothelial, stromal and immune compartments, and further organized into 18 epithelial-dominated, cancer associated fibroblast-dominated, and immune-rich spatial niches. Within the epithelial compartment, we detected an ERG⁺p53⁺ luminal population whose abundance is independently associated with poor overall and progression-free survival. In the stroma, we defined extracellular matrix remodeling-related cancer associated fibroblast and smooth muscle cell lineages, including a periglandular CD105^high^ niche with strong stromal-immune connectivity that is selectively associated with worse clinical outcome. Finally, cumulative immune niche burden correlated with histological inflammation and stratifies for worse patient survival. Together, these data provide a spatially resolved single-cell atlas of primary PCa and reveal stromal-immune-epithelial niches with prognostic relevance beyond Gleason grade.

## Introduction

Prostate cancer (PCa) displays extensive intra-tumoral and inter-patient heterogeneity, ranging from indolent to clinically progressive tumors despite definitive local therapy^1–6^. Current clinical risk stratification relies heavily on histopathological Gleason grading; however, intermediate Gleason grade categories frequently fail to capture the biological diversity and prognostic variability observed among patients^7,8^. This limitation contributes to both over- and undertreatment and underscores the need for improved biomarkers that more accurately reflect tumor biology and predict clinical outcomes.

Increasing evidence suggests that this shortcoming arises partly from the complex and spatially organized architecture of the tumor microenvironment (TME), not fully captured on standard diagnostic slides^9–11^. At the epithelial level, PCa typically arises from luminal epithelial cells within glands that canonically harbor basal and luminal lineages interspersed with rare neuroendocrine (NE) cells^12,13^. Single cell atlases of the prostate and proximal urethra have expanded the epithelial repertoire to include club-like and hillock-like cells, which are linked to epithelial homeostasis and injury responsive programs and can emerge under inflammatory conditions^14–16^. These noncanonical states illustrate the marked epithelial plasticity of the prostate, enabling transitions toward androgen receptor (AR) independent phenotypes, including treatment emergent neuroendocrine prostate cancer and double negative castration resistant PCa, often in the context of TP53/RB1 perturbations and alternative lineage regulators^17,18^.

Although traditionally considered “immune cold,” localized PCa contains a structured and functionally constrained immune microenvironment. Spatial and single cell analyses show that immune evasion reflects not only limited infiltration but also spatially restricted suppression, including exhausted T cells juxtaposed with suppressive myeloid populations and stromal programs that bias cytokine signaling^4,16,19^. Moreover, immune-epithelial interactions involving club-like epithelial cells, immunosuppressive myeloid populations, and androgen receptor (AR)-positive stromal cells define coordinated multicellular hubs within the tumor microenvironment that are increasingly recognized as clinically relevant^16,20–26^.

The stromal compartment, comprising smooth muscle cells (SMCs), cancer associated fibroblasts (CAFs), pericytes, and extracellular matrix (ECM), represents a major yet incompletely characterized component of the PCa TME. Single-cell transcriptomics has outlined diverse CAF states with myofibroblastic, inflammatory, and antigen-presenting features, and has linked stromal programs to angiogenesis, immune modulation, bone metastasis and androgen deprivation therapy (ADT) resistance^27–31^. However, the spatial organization, lineage relationships, and clinical relevance of these stromal phenotypes, and their interactions with immune and epithelial compartments, remain poorly resolved. Spatially resolved single-cell proteomic technologies, such as imaging mass cytometry (IMC)^32^, now enable multiplexed in situ mapping of the TME at cellular resolution. While IMC studies have provided insights into CAF biology in other tumor types^33,34^, a comparable spatial proteomic atlas of primary PCa is lacking.

Here, we construct a spatially resolved single-cell atlas of primary PCa using a prostate tissue-tailored 34-plex IMC panel applied to a large, clinically annotated radical prostatectomy tissue microarray (195 patients, 523 regions, 2.19 million cells). We resolve epithelial, stromal, endothelial, and immune cellular populations and their organization into compartments, define CAF/SMC heterogeneity, and identify 18 recurrent spatial niches, that form hierarchical epithelial-stromal-immune ecosystems. We further evaluate the clinical relevance of these architectures, including a CD105⁺ periglandular CAF niche, that extend risk stratification beyond conventional histopathology.

## Results

### Spatial proteomics defines the complex and heterogeneous architecture of primary PCa TME

We performed a comprehensive spatial proteomic analysis using imaging mass cytometry (IMC) on a tissue microarray (TMA) cohort of 195 primary PCa specimens. Quality control filtering removed regions with staining artifacts and segmentation errors, resulting in a final dataset of 523 high-quality ROIs and 2.19 million single cells for downstream analysis. This approach enabled high-dimensional, spatially resolved single-cell analysis of epithelial, stromal, immune, and endothelial compartments within intact tissue architecture, including quantitative compositional analysis of cell populations, and spatial organization of niches (Fig. 1a). The cohort includes treatment-naive primary tumors collected at radical prostatectomy (RP) as part of the European Multicenter Prostate Cancer Clinical and Translational research group (EMPaCT)^35,36^ and is supported by comprehensive clinical and pathological annotations including patient-level Gleason grade group, and longitudinal clinical endpoints such as recurrence, progression, and overall survival (Methods, Supplementary Fig. 1a). The TMA includes up to four cores per patient, capturing the multifocal tissue architecture and intra-patient heterogeneity of PCa. Histopathological evaluation for core-level Gleason group, presence of reactive stroma (stromogenic features), and inflammation status was performed by an expert pathologist on serial sections stained with hematoxylin and eosin (H&E).

**Figure 1:**
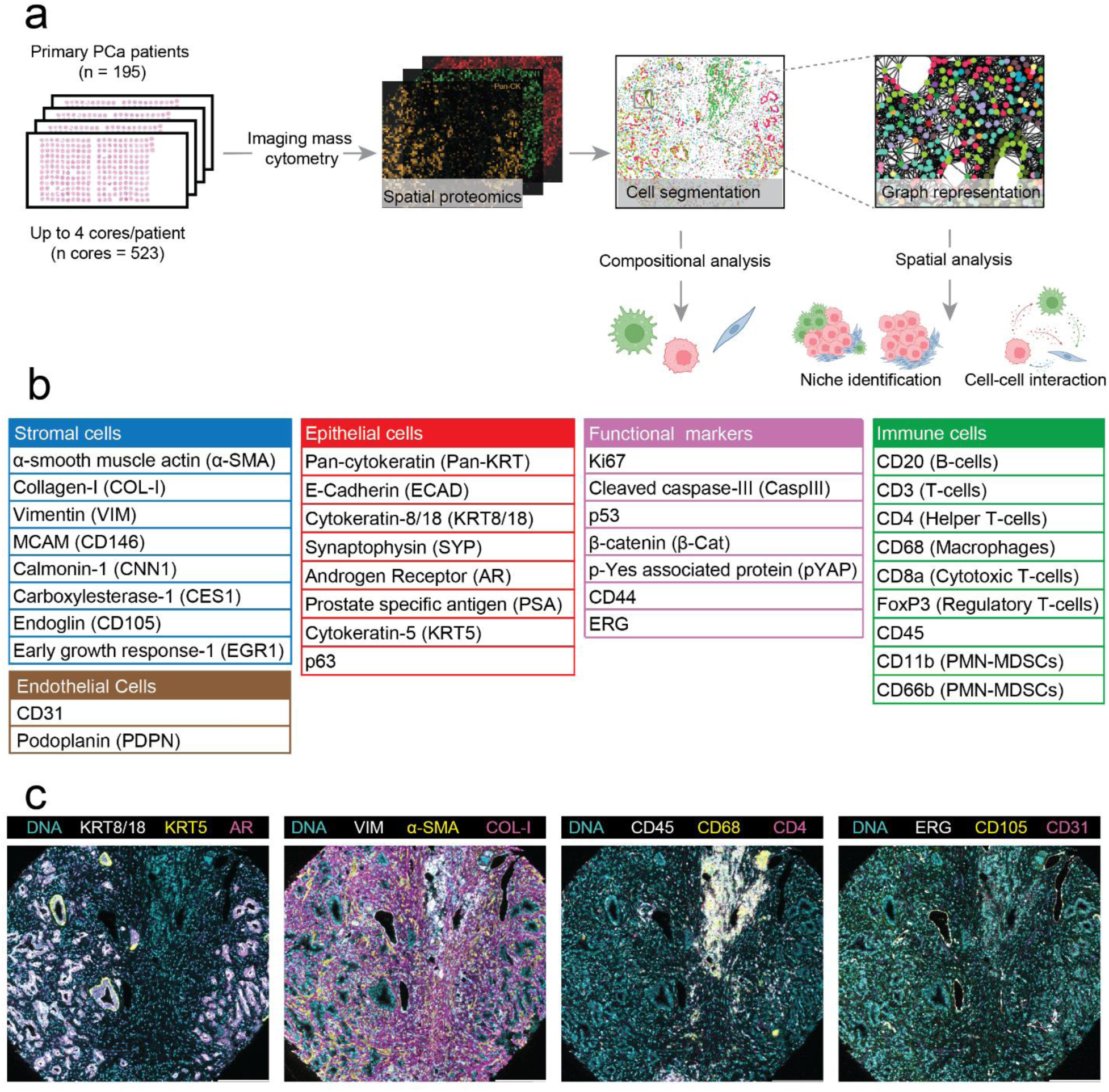
Imaging mass cytometry (IMC) workflow and marker panel for profiling the prostate cancer (PCa) tumor microenvironment (TME). **a** Overview of the study workflow, from primary PCa tissue microarrays (TMAs) through IMC acquisition to downstream data processing, including segmentation and analyses of cellular composition and spatial organization (schematic created with BioRender.com). **b** Composition of the 34-plex IMC antibody panel. Markers were selected to identify major cellular compartments (epithelial, immune, endothelial, cancer-associated fibroblasts (CAFs), and smooth muscle cells (SMCs)) and to quantify functional states, including proliferation, apoptosis, oncogenic and tumor-suppressor pathways. Iridium-191/193 DNA intercalator was included to delineate nuclei. **c** Representative IMC images illustrating simultaneous visualization of the major tissue compartments (epithelial, stromal, immune, and endothelial). Marker and DNA colors are pseudo-colored for visualization and do not reflect the metal isotope channels. Scale bar: 200 ∼μm.

Histopathological review of matched H&E sections confirmed the expected multifocal architecture of PCa and revealed marked differences with respect to Gleason grade across cores from the same patient (Supplementary Fig. 1b). In 72.9% of patients, at least one of all cores displayed a Gleason pattern that differed from the patient-level Gleason grade group, including cases where high-grade foci occurred in tumors clinically classified as low grade (Supplementary Fig. 1b). This finding underscores the epiphenomenon of intratumoral heterogeneity of primary PCa and further highlights the limitations of conventional Gleason grading in capturing clinically relevant disease biology, thereby motivating the need for spatially resolved molecular characterization of the PCa TME^37^.

Using a 34-plex IMC antibody panel, we resolved the major cellular compartments of the PCa TME, including epithelial, stromal, immune, and endothelial populations (Fig. 1b). The IMC panel was designed to capture both canonical cell lineages and functional states, with particular emphasis on resolving stromal heterogeneity (SMCs, pericytes, and diverse CAF states, including inflammatory and matrix-remodeling programs). CAF and SMC marker selection was informed by recent single-cell transcriptomic analysis of primary PCa^38^, with carboxylesterase 1 (CES1) and CD105 selected as inflammatory (iCAF) and myofibroblast CAF (myCAF) markers, respectively. YAP1 phosphorylated at tyrosine 357 (pYAP) was selected as a marker of CAF activation, reflecting the established role of YAP signaling in driving fibroblast reprogramming to matrix-producing myCAF phenotypes^39,40^.

Qualitative assessment of IMC images confirmed the specificity and spatial organization of marker staining across tissue compartments (Fig. 1c). The expected glandular architecture was preserved, with epithelial cells surrounded by stromal and immune elements. Notably, AR staining was abundant but not limited to epithelial luminal cells: stromal cells also displayed nuclear AR expression (Fig. 1c), consistent with previous publications^41–43^.

### Single-cell phenotyping reveals diverse epithelial, immune and stromal subpopulations

To systematically define cellular populations within the PCa TME, we performed single-nuclei segmentation and quality control followed by unsupervised clustering and marker-based cell annotation (Fig. 2a). This yielded a total of 2’191’967 high-quality single-cell profiles across all patients, classified into 34 cell types. The epithelial compartment represented the largest fraction (48%), followed by stromal (30.3%), immune (15.1%), and endothelial compartments (3.7%), while a small fraction of cells (3%) remained unclassified and was excluded from the analysis. UMAP visualization demonstrated a clear separation between epithelial and stromal compartments (Fig. 2b), with cells from different patients largely intermixing across clusters. Only a small number of rare subclusters appeared to be patient-specific, consistent with biological heterogeneity rather than batch effects. Within the epithelial compartment, 9 phenotypically distinct populations were identified, including luminal (KRT8/18^+^, AR^+^) and basal (KRT5^+^, p63^+^) epithelial cells, and a rare mixed cluster with both luminal and basal markers (KRT8/18^+^ KRT5^+^), suggestive of intermediate or transient states. Additional subpopulations captured distinct proliferative and oncogenic programs, including luminal Ki67^+^ proliferative cells, luminal ERG^+^ tumor cells consistent with TMPRSS2-ERG fusion, and luminal p53-expressing cells (Fig. 2a; Supplementary Fig. 2a). Notably, a small subset of ERG^+^ cells (6.1%) also expressed p53, potentially reflecting tumor cells under genotoxic stress or enriched for TP53 mutations. Neuroendocrine (NE) cells identified by synaptophysin expression constituted one of the smallest populations, consistent with recent studies implicating NE plasticity during disease progression and therapeutic resistance^18,44^.

**Figure 2.**
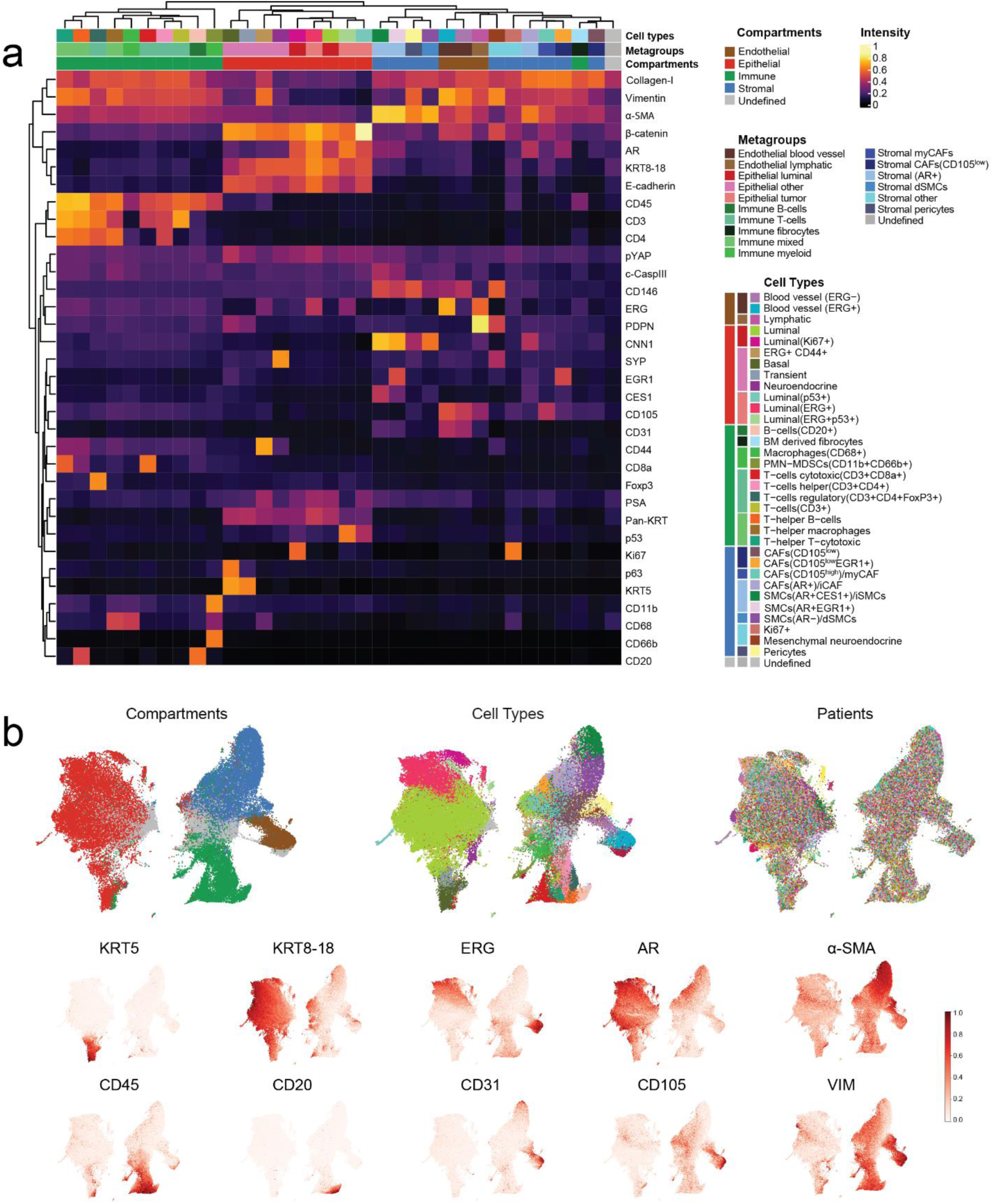
Single-cell phenotyping resolves diverse cellular populations in the PCa tumor microenvironment. **a** Clustered heatmap of normalized mean marker expression across the 34 annotated cell types delineates major cellular compartments and highlights distinct protein expression signatures across epithelial, stromal, immune, and endothelial populations. **b** UMAP embedding of all cells (n = 2’191’967) colored by compartment, cell types and patients (top row) and specific markers (bottom two rows).

The immune compartment comprised 11 diverse lymphoid and myeloid populations, including T-helper cells (CD3^+^CD4^+^), cytotoxic (CD3^+^CD8a^+^), and regulatory T cells (CD3^+^CD4^+^FOXP3^+^), macrophages (CD68^+^), B cells (CD20^+^) and polymorphonuclear myeloid-derived suppressor cells (PMN-MDSCs) (CD11b^+^ and CD66b^+^), a population previously reported to contribute to castration resistance (Fig. 2b, Supplementary Fig. 2b)^45^. Among CD45^+^ cells, we identified a distinct population co-expressing general stromal markers (vimentin, α-SMA, and collagen-I) (Fig. 2a), consistent with previously reported fibrocytes, cells of hematopoietic origin expressing low CD45^+^ levels along with typical fibroblast markers^46,47^. Fibrocytes have been recently associated with adverse prognosis and antigen presenting (apCAF)-like states^48^, suggesting a potential role in stromal-immune crosstalk. Endothelial cells (CD31^+^ CD105^+^) were further stratified based on ERG expression (Fig. 2a, Supplementary Fig. 2c). Given the role of ERG in regulating endothelial cell identity and homeostasis, and its downregulation under inflammatory conditions^49^, ERG^+^ and ERG^-^vessels may reflect quiescent, versus activated, inflammatory angiogenic states, respectively. Lymphatic endothelial cells were distinguished from blood endothelial vessels by PDPN^+^ expression (Supplementary Fig. 2c).

### Identification of distinct CAF populations with ECM-remodeling and SMC-like properties

We next focused on resolving fibroblast heterogeneity within the non-immune stroma. Unsupervised clustering of stromal cells identified 10 distinct CAF populations, reflecting diverse phenotypic and functional states (Fig. 2a, 3a-b). Most stromal cells separated into two major phenotypic groups: a CAF group with expression patterns typical of ECM-producing cells (VIM^+^, COL-I^+^and *α*-SMA^low^ expression), and a second group displaying SMC-like features (CNN1^+^, CD146^+^, *α*-SMA^high^) (Fig. 3a-b, d).

**Figure 3.**
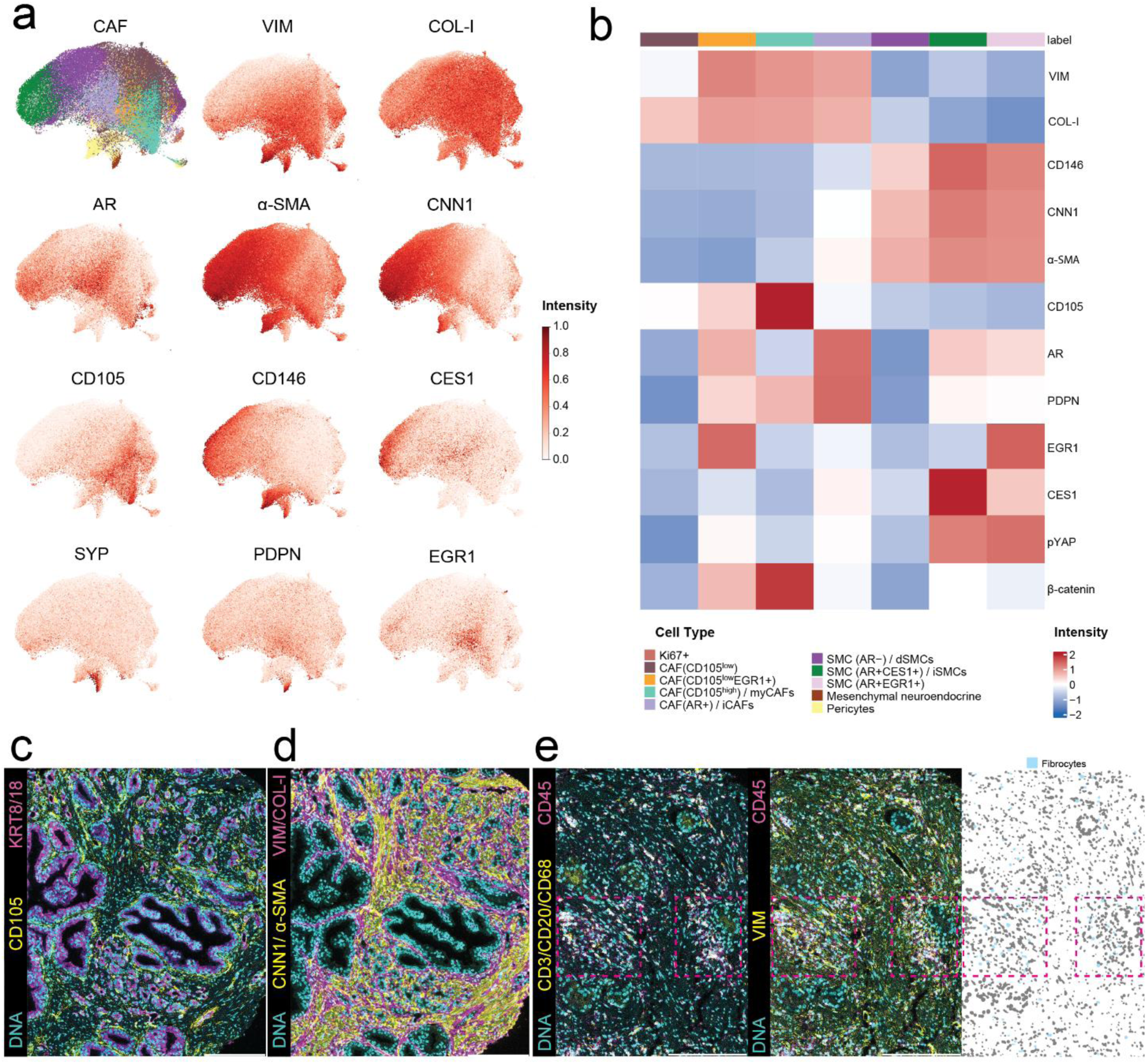
Identification of CAF phenotypes. **a** UMAP of all CAF cells across the cohort, computed using CAF-relevant markers (*α*-SMA, vimentin, collagen I, CD146, CNN1, CD105, AR, EGR1, and CES1). Top left: cells colored by CAF subcluster. **b** Heatmap of mean marker expression per CAF subcluster, highlighting subtype-specific expression patterns and putative functional differences. Values represent average z-scored marker expression per cluster. **c-e** Representative IMC regions of interest (ROIs) illustrating the spatial localization and morphology of selected CAF phenotypes: periglandular enrichment of myCAFs (**c**), distinct phenotypic features of ECM-producing myCAF-cells (VIM/Collagen-I) and SMC cells (*α*-SMA/CNN1/CD146) (**d**), and identification of fibrocyte-like cells co-expressing CD45 but negative for the rest of immune markers (CD20, CD3, CD68) (left), with VIM (middle) and with the cell-mask indicated (right) (**e**). Scale bar: 200 ∼*μ*m.

The phenotypic signature of the CAF group, combining high matrix protein expression (COL1A1) with lower contractile features (*α*-SMA, CNN1), is consistent with ECM-producing, myCAF-like cell phenotype. Within CAFs, further heterogeneous subpopulations were defined by differential expression of CD105, AR, and EGR1, giving rise to CD105^high^, CD105^low^, CD105^low^EGR1^+^ and AR^+^ populations. CD105, an auxiliary TGF-*β* receptor, has been associated with myCAFs in late-stage PCa by scRNA-seq analysis^38^, therefore CD105^high^ are annotated as myCAFs. Notably, CD105^high^ myCAFs displayed a preferential periglandular localization, suggesting a role in tumor-stroma crosstalk and ECM remodeling (Fig. 3c). The CD105^low^ populations were previously found in higher abundance in non-tumor, prostate tissues (scRNASeq) compared to PCa^38^. The expression of EGR1, a transcription factor expressed in fibroblasts in response to injury and stress cues^50^ and upstream activator of profibrotic TGFβ^51^, is found in a subset of CD105^low^ cells. The CD105^low^ and CD105^low^EGR1^+^ possibly represent distinct activation/ intermediate substates of CAFs, for which their phenotype remains to be characterized. PDPN expression was found in higher levels in CAF subpopulations compared to SMCs (Fig. 3b), in line with myCAF/CAF annotation in breast cancer^52,53^. The AR+ CAF subpopulation expresses moderate levels of CES1, a known inflammatory-associated CAF marker, and lacks myCAF marker expression (CD105) which were found to be AR negative in our previous findings^39^. Thus, the AR+CES1+ CAF subpopulation is annotated as iCAF.

The SMC cluster retains contractile features consistent with a key role of the prostate stroma in maintaining structural rigidity of the gland (Fig. 3b) and could be further subdivided into three types: AR^+^CES1^+^, AR^+^EGR1^+^, and AR^-^. CES1^+^ stromal cells were previously identified in SMC of the benign prostatic stroma and associated with favorable prognosis^54^. Consistent with prior reports^41^, stromal AR expression showed marked heterogeneity, with AR^+^ populations present in both CAF and SMC compartments, while loss of AR in SMCs is associated with PCa^25^.

The SMC subpopulation, marked as AR^+^CES1^+^, is similar to intact, interstitial, non-activated SMCs (Damisch et al dataset, M3)^38^ since also CES1^+^ is rarely found in vascular SMCs. The AR^+^CES1^+^ is thus labeled as interstitial SMCs (iSMCs).

The profile of AR^+^EGR1^+^ subpopulation with moderate CES1 expression, resemble a mix of interstitial non-tumor associated SMCs (non-dedifferentiated) and vascular SMCs, similar to the Damisch et al., M1/M2/M3 clusters of mural cells^38^.

The AR^-^ SMC subpopulation is indicative of interstitial, tumor-associated SMCs based on lower levels of CD146 and contractile markers (CNN1, aSMA) compared to the other SMC subpopulations, as previously found in interstitial SMCs undergoing tumor-associated dedifferentiation^38^. Therefore, AR^-^ SMCs are annotated as dedifferentiating SMCs (dSMCs). Overall, these findings indicate that stromal AR characterization could be further defined from general (ECM-remodeling or SMC-rich stroma) to CAF/SMC subtype-specific and based on association with specific markers (CES1, EGR1, CD105), towards interrogation of potential prognostic stratification.

In addition, we identified shared and rare stromal populations independent of CAFs/SMCs classification. These include a rare (<0.2%) proliferative stromal subset marked by Ki-67, stromal pericytes (CD146^+^, *α*-SMA^+^, CNN1^-^) with characteristic perivascular localization, and a rare (<0.5%), islet-like population expressing PDPN, CD146, and the neuroendocrine marker SYP. Finally, qualitative assessment confirmed the lack of immune markers in the CD45^low^ expressing fibrocytes (Fig. 3e).

Together, these results reveal a highly complex stromal landscape in primary PCa, characterized by multiple CAF states with distinct structural, signaling, and potential functional properties, as well as diverse patterns of AR expression across stromal subpopulations.

### Cell type composition and association with clinical outcome

To investigate whether variation in cell-type composition relates to clinical outcome, we aggregated cell-type frequencies at a patient-level and performed hierarchical clustering, revealing six patient groups (P1-P6) with distinct cell type compositions (Fig. 4a-b). The two largest clusters (P3 and P5) were characterized by relative enrichment for luminal epithelial and tumor epithelial cells, respectively; P2 and P4 were enriched for diverse CAF and SMC populations, whereas P6 was predominantly enriched for immune populations (Fig. 4a-b). To assess whether the intra-patient heterogeneity previously observed across Gleason grade groups is also reflected at the level of cellular compositions, we independently clustered core-level cell type distributions, identifying 6 core-level groups (C1-C6) (Supplementary Fig. 3a). Consistent with the patient-level analysis, C2 and C5 were dominated by luminal epithelial and tumor epithelial cells, respectively. We next evaluated the correspondence between core-level and patient-level clusters. We observed pronounced intra-patient heterogeneity, with cores from the same patient frequently assigned to different compositional clusters (Supplementary Fig. 3b). However, C2 and C5 appeared largely mutually exclusive: patients with at least one core in C2 did not tend to harbor cores in C5, and vice versa, suggesting the presence of distinct, patient-specific epithelial dominant programs rather than co-existing compositional states.

**Figure 4.**
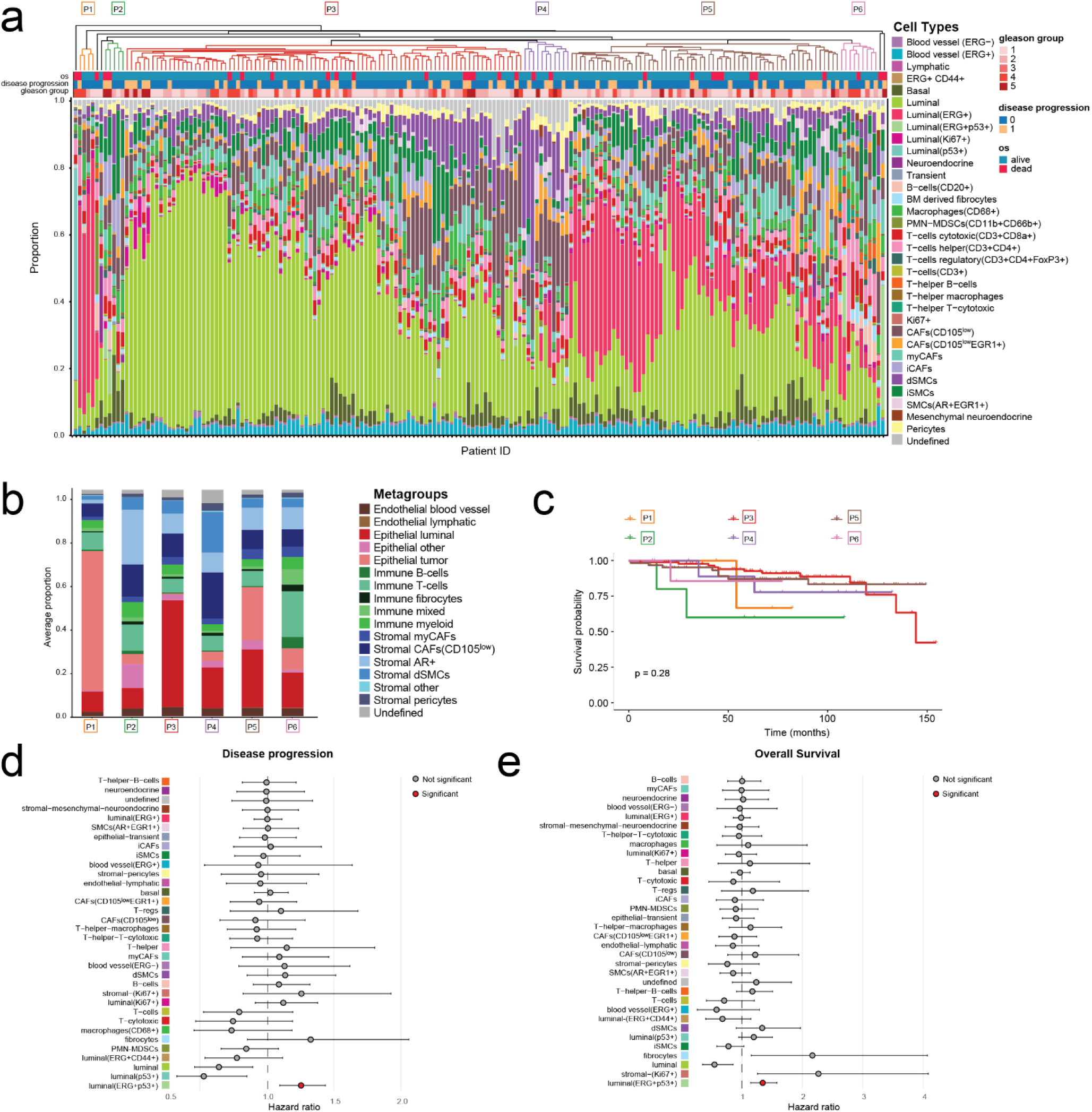
Cell type composition and association with clinical outcomes. **a** Hierarchical clustering of patient-level cell-type proportions based on the full cell-type annotation resolves six patient cluster groups (P1-P6), indicated by colored branches in the dendrogram. **b** Mean cellular metagroup distribution across the six patient clusters. **c** Kaplan-Meier survival analyses stratified by patient-level group; statistical significance was assessed using a two-sided log-rank test. **d-e** Cox proportional hazards models assessing associations between cell-type proportions and overall survival (**d**) or disease progression (**e**). Hazard ratios with 95% confidence intervals (horizontal lines) are shown; *p*-values were adjusted for multiple testing using the Benjamini-Hochberg false discovery rate (FDR) procedure; statistical significance was defined as FDR-adjusted *p* < 0.05.

We then assessed whether these patient compositional groups stratified clinical outcome. Kaplan-Meier analysis revealed no statistically significant association between these groups and overall or progression-free survival probability (Fig. 4c, Supplementary Fig 3c). We next evaluated associations between individual cell types and clinical outcomes using Cox proportional hazards models based on patient-level cell-type frequencies (Fig. 4d-e). Among all populations tested, the luminal ERG^+^p53^+^ epithelial population emerged as the only cell type significantly associated with both overall and progression-free survival, with higher abundance linked to worse outcomes. In contrast, the luminal ERG^+^ and p53^+^ single-positive populations were not associated with any clinical endpoint. Although TP53 alterations occur at low frequency in primary PCa (<5%) relative to their markedly higher frequency at the metastatic stage (40-60%)^55,56^, our data suggest that co-occurrence of ERG and p53 expression defines a particularly aggressive epithelial state, already present in localized tumors. Finally, fibrocytes showed a trend toward association with both overall survival and progression-free survival, but this did not remain significant after multiple testing correction. This trend is nonetheless consistent with recent spatial single-cell studies describing apCAF-like niches associated with immune-rich microenvironments and adverse clinical features^48^.

### Spatial niche analysis reveals hierarchical tumor-stroma organization in primary PCa

To characterize spatial co-occurrence patterns among cell types, we performed niche identification analysis as previously described^57^. This approach identified 18 distinct cellular niches with heterogeneous composition, area occupancy, and frequency across cores (Fig. 5a), as well as defined patterns of co-occurrence and mutual exclusion (Fig. 5b). Unsupervised hierarchical clustering of niche cell type enrichment stratified these into three major categories (Fig. 5a): (i) epithelium-enriched niches (niches 1-8), (ii) CAF/SMC dominant niches with low gradients of epithelial or immune cells, (niches 9-15) and (iii) immune-enriched niches (niches 16-18).

**Figure 5.**
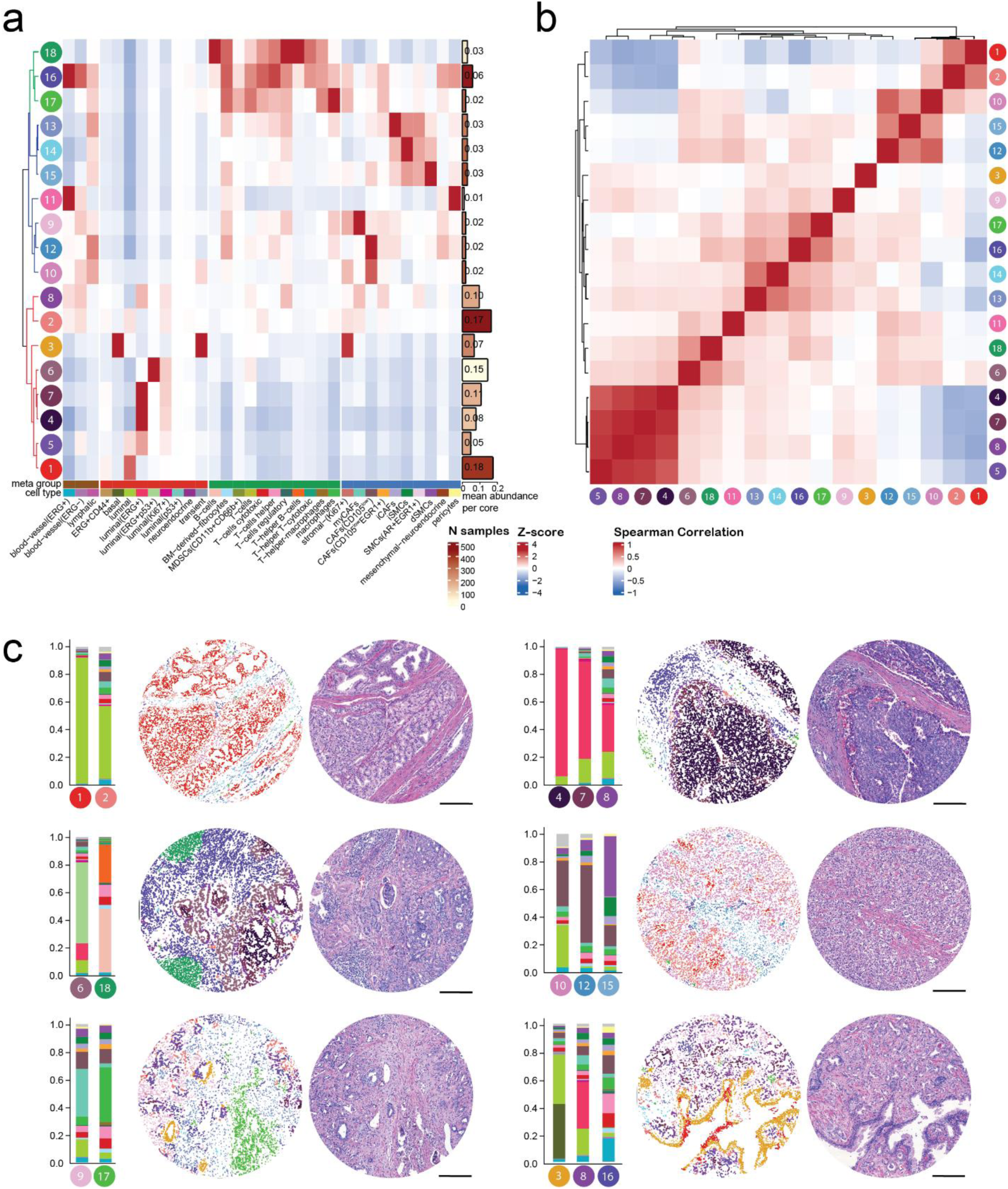
Spatial distribution of cell types organized in cellular niches. **a** Heatmap of z-scored mean cell type composition across 18 identified niches, with hierarchical clustering stratifying niches into three distinct categories: epithelial-dominated (1-8), CAF-dominant (9-15), and immune-enriched (16-18). Columns represent cell types, and rows represent niches, with the barplot on the right indicating mean niche abundance per core (bar length) and frequency in the cohort (bar color) of each niche. **b** Clustered pairwise Spearman correlation matrix of niche abundance across samples, highlighting co-occurrence (red) and mutual exclusion (blue) patterns between all identified niches. **c** Representative IMC cores with their corresponding H&E images, illustrating the spatial architecture of selected niches, including luminal epithelium, tumor core, tumor-stroma interface, and CAF- and immune-enriched regions. Adjacent stacked barplots show the mean cell type composition of the dominant niches in these cores (containing at least 15 cells of the niche). Scale bar: 200 ∼μm.

The epithelium-enriched niches were divided into luminal (niches 1, 2, 3) and tumor-associated groups (niches 4-8). Niches 1 and 2 were the most prevalent and occupied the largest areas within tissue cores. Niche 1 represents a near-pure luminal compartment (>90% luminal cells), whereas niche 2 comprised luminal cells interspersed with mixed stromal populations, including CAFs, SMCs, endothelial cells, and sparse immune infiltrates. These two niches frequently co-occurred in the same cores (Spearman r=0.62) (Fig. 5b), with niche 2 forming either a structured interface between niche 1 and the surrounding TME, or a less differentiated epithelial state with extensive stromal infiltration (Fig. 5c, Supplementary Fig. 4a-b). Niche 3 corresponded to canonical glandular structures, comprising luminal and basal layers (Supplementary Fig. 4a-b), and enriched for transient epithelial cells.

Tumor-associated niches (niches 4-8) defined by ERG expression exhibited a hierarchical spatial organization. Niche 4 constituted of ERG^+^ tumor cells, localized to inner tumor cores, whereas niche 5 contained a mixture of ERG^-^ and ERG^+^ epithelial populations. These were surrounded by niche 7, representing a layer of tumor-stroma interaction zones, and further by niche 8, a complex interface with ERG tumor cells and enriched for immune cells (primarily T cells and macrophages) and CAF subtypes (including myCAF and CAF CD105^low^) (Fig. 5c). Tumor-associated niches formed a cohesive spatial module, with strong co-occurrence (mean Spearman r=0.89), reflecting a consistent core-to-margin organization (Fig. 5b). In contrast, they were negatively correlated with luminal niches 1 and 2, indicating mutually exclusive epithelial compartments. Finally, a distinct niche enriched for ERG^+^p53^+^ tumor cells and proliferative luminal populations (niche 6), defined a rare but spatially extensive context when present, suggesting localized expansion of aggressive epithelial states.

Non-immune stroma dominant niches were segregated into CAF (niches 9, 10, 12) or SMC enriched groups (niches 13-15). Niches 9 and 10 localized at epithelial-stromal boundaries occupying a small area of the core (∼2%); niche 9 is enriched in myCAF cells, and recapitulated the peri-glandular CD105^high^ pattern observed in IMC images (Fig. 5c); it showed inconsistent presence across samples, suggesting context-dependent microenvironmental cues. Conversely, niche 10 was enriched in CAF CD105^low^ cells and was more consistently detected across samples. These niches were mutually exclusive, suggesting that CD105 presence defines functionally distinct epithelial-stromal boundary regions. Niche 12, while similarly enriched as niche 10 for CAF CD105^low^ cells, lacked epithelial cells and localized exclusively within the stromal compartment.

Niches 13-15 comprised mixed CAF and SMC populations together with 15–20% immune cells, but no epithelial cells (Supplementary Fig. 4b). Each of these heterogeneous niches was dominated by distinct stroma subtypes: iCAFs in niche 13, iSMCs in niche 14, and dSMCs in niche 15. These niches exhibited preferential co-occurrence patterns: niches 13 and 14 clustered together, while niches 10, 12, and 15 formed a separate group (Fig. 5b). Interestingly, niche 13, enriched for iCAF population, occasionally co-occurred with immune-enriched niches 16 and 17 (Fig. 5b), consistent with coordinated spatial patterning of inflammatory stroma and immune compartments, and in line with the reported iCAF role of CES1^+^ cells^38,39^.

Immune-enriched niches (niches 16-18) displayed diverse compositions. Niche 16 represented a mixed immune microenvironment, comprising roughly equal amounts of endothelial cells, CD4^+^ T-helper cells, CD8^+^ cytotoxic T cells, macrophages, and a minor fraction of CAF populations. Niche 17 was dominated by macrophages (∼40%) including the rarer PMN-MDSC subpopulation. Niche 18 consists of B cells and T cells in high proximity, resembling a tertiary lymphoid structure (TLS), suggesting adaptive immune responses against tumor cells^58,59^. Vascular structures, consisting of an inner endothelial layer surrounded by pericytes, were captured as a distinct niche 11 (Supplementary Fig. 4a). Collectively, these data reveal a hierarchical and spatially coordinated organization of epithelial, stromal, and immune compartments, in which distinct CAF and immune niches define specialized microenvironmental interfaces.

### Niche abundance associations with clinical outcomes

To determine whether spatial niche organization is associated with clinical and histopathological features, we quantified niche abundance per core and performed unsupervised clustering, yielding five clusters of tissue cores with distinct niche composition profiles (Fig. 6a). Cluster 1 was enriched for luminal niches 1 and 2, whereas cluster 2 was defined by ERG^+^ tumor-associated niches (niches 4, 5, 7 and 8), consistent with previous observations. Cluster 3, the smallest of all, was composed predominantly of niche 6, enriched for ERG^+^p53^+^ tumor cells. Finally, clusters 4 and 5 displayed mixed compositions, combining luminal niches with the immune-driven and CAF-driven niches, respectively. These niche-based clusters were not visually associated with clinical parameters.

**Figure 6.**
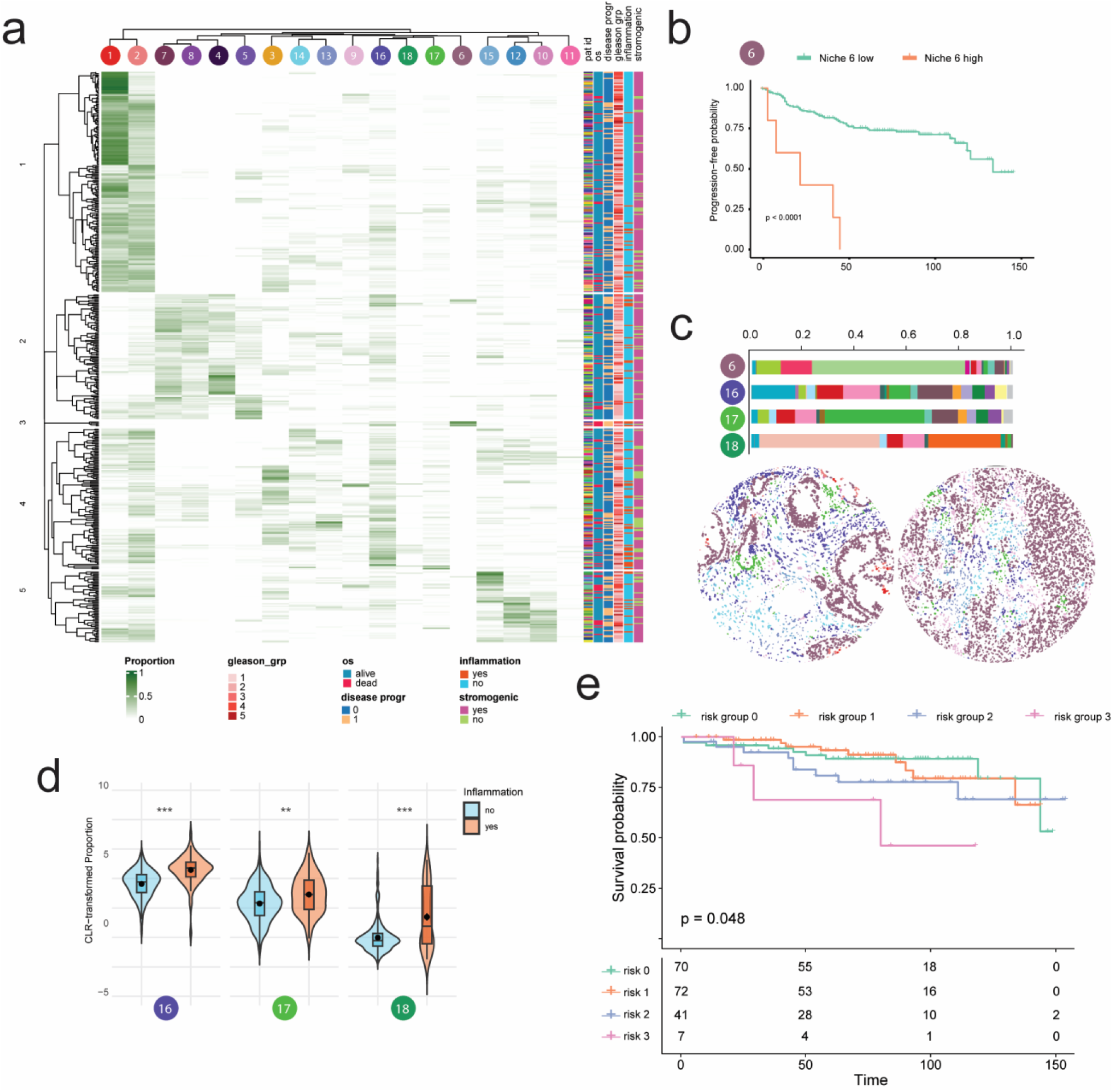
Niche abundance and clinical associations. **a** Clustered heatmap of niche abundance scores per tumor core, with associated metadata shown on the right. **b** Kaplan-Meier survival analysis, with patients grouped into high and low categories based on niche 6 abundance, defined at the core level using the cohort median and aggregated to the patient level (high if ≥1 high-abundance core); p-values were computed using a two-sided log-rank test. **c** Stacked barplot of mean cell type composition across cores of niches 6, 16, 17 and 18 (top), with two example cores (bottom). **d** Distribution of niche abundance per inflammation status, indicating enrichment of immune-associated niches in inflamed ROIs. Statistical significance was assessed using a Wilcoxon rank test after CLR-transformed niche proportion values, and p-values were adjusted for multiple testing using a Benjamini-Hochberg correction. **e** Kaplan-Meier survival analysis stratified by immune niche-derived risk score. Patients were stratified into four risk groups based on high abundance (>75^th^ percentile of abundance per niche across all cores) of 0, 1, 2 or all 3 immune-associated niches 16-18.

We next assessed the prognostic relevance of individual niches. Kaplan-Meier analysis demonstrated that high abundance of niche 6 was associated with reduced overall survival (Fig. 6b), consistent with the aggressive phenotype and adverse prognostic value of the ERG^+^p53^+^ cell type previously identified at the single-cell level (Fig. 4d-e). Importantly, the sole presence of ERG^+^p53^+^ niche was not associated with Gleason grade group (Supplementary Fig. 5a), indicating that spatially resolved features capture clinically relevant information beyond standard histopathological stratification. Notably, qualitative inspection revealed that niche 6 frequently co-occurred with immune-related niches 16, 17, and 18 within the same tissue cores (Fig. 6c, Supplementary Fig. 5a).

To formally assess the relationship between spatial niches and inflammation, we compared the cumulative abundance of immune-associated niches across inflamed versus non-inflamed cores. All three immune niches (16-18) were significantly enriched in inflamed cores (Fig. 6d), supporting the validity of the spatial niche classification and its ability to capture clinically relevant inflammatory states. Finally, we derived an immune niche-based risk score by stratifying patients in four risk groups, depending on whether there is a strong presence (> 75^th^ percentile) of 0, 1, 2 or all 3 immune niches in at least one out of the patient cores. Increased representation of immune niches was associated with reduced lower overall survival (Fig. 6e), indicating that cumulative niche burden rather than individual niche presence captures the prognostically relevant inflammatory microenvironment. This finding captures and refines the signal from histological inflammation annotations, which were likewise significantly associated with worse overall survival (Supplementary Fig. 5b).

Together, these findings demonstrate that spatial niche composition captures both tumor-intrinsic aggressive states and coordinated stromal–immune organization, providing clinically relevant information beyond conventional pathological features.

### CD105^high^ stromogenic niche associates with clinical outcome and mediates stromal–immune interactions

Stromal remodeling and the emergence of a “reactive” or stromogenic microenvironment have been repeatedly linked to adverse prognosis in PCa, supporting a prognostic role of stromal patterning beyond epithelial features alone^11,60,61^. In our cohort, Kaplan-Meier analysis did not reveal a significant association between histologically defined stromogenic cores and progression-free survival (*p*=0.57, Supplementary Fig. 6a), highlighting the limitations of binary histopathological classification without relative quantitative assessment of stromal composition^60,61^.

To refine this analysis, we examined the distribution of stromal-associated niches across stromogenic versus non-stromogenic cores (Fig. 7a). This revealed different enrichment patterns, with niches enriched in CAF stromal cells (niches 2 and 8-10) over-represented in stromogenic cores, except for niche 13 (iCAF rich) and niche 12 (CD105^low^). Whereas niches with SMCs-rich stroma (niches 14-15) were found under-represented in stromogenic cores, indicating distinct stromal programs underlying similar histopathological appearances. Among these, niche 9, characterized by a pronounced enrichment of myCAF cells, showed a significant association with reduced progression-free survival (*p*=0.0182; Fig. 7b). To determine whether this effect was driven by the myCAF cell population alone, we compared niche-level and cell type–level associations. In the previous cell-type-focused Cox analysis (Fig. 4d-e), the abundance of myCAFs was not independently associated with outcome. Consistently, Kaplan-Meier analysis based solely on myCAFs abundance confirmed the lack of association with progression-free survival (*p*=0.24; Fig. 7c). Although myCAF cells were also enriched in niches 2 and 8 (Fig. 5a), both of which were overrepresented in stromogenic cores (Fig. 7a), neither niche was associated with progression-free survival (*p*=0.68 and *p*=0.68 respectively; Supplementary Fig. 6b-c). Together, these findings indicate that neither stromogenic histology nor myCAF abundance alone is sufficient to explain clinical outcome, underscoring the importance of spatial context in defining functionally relevant stromal states.

**Figure 7.**
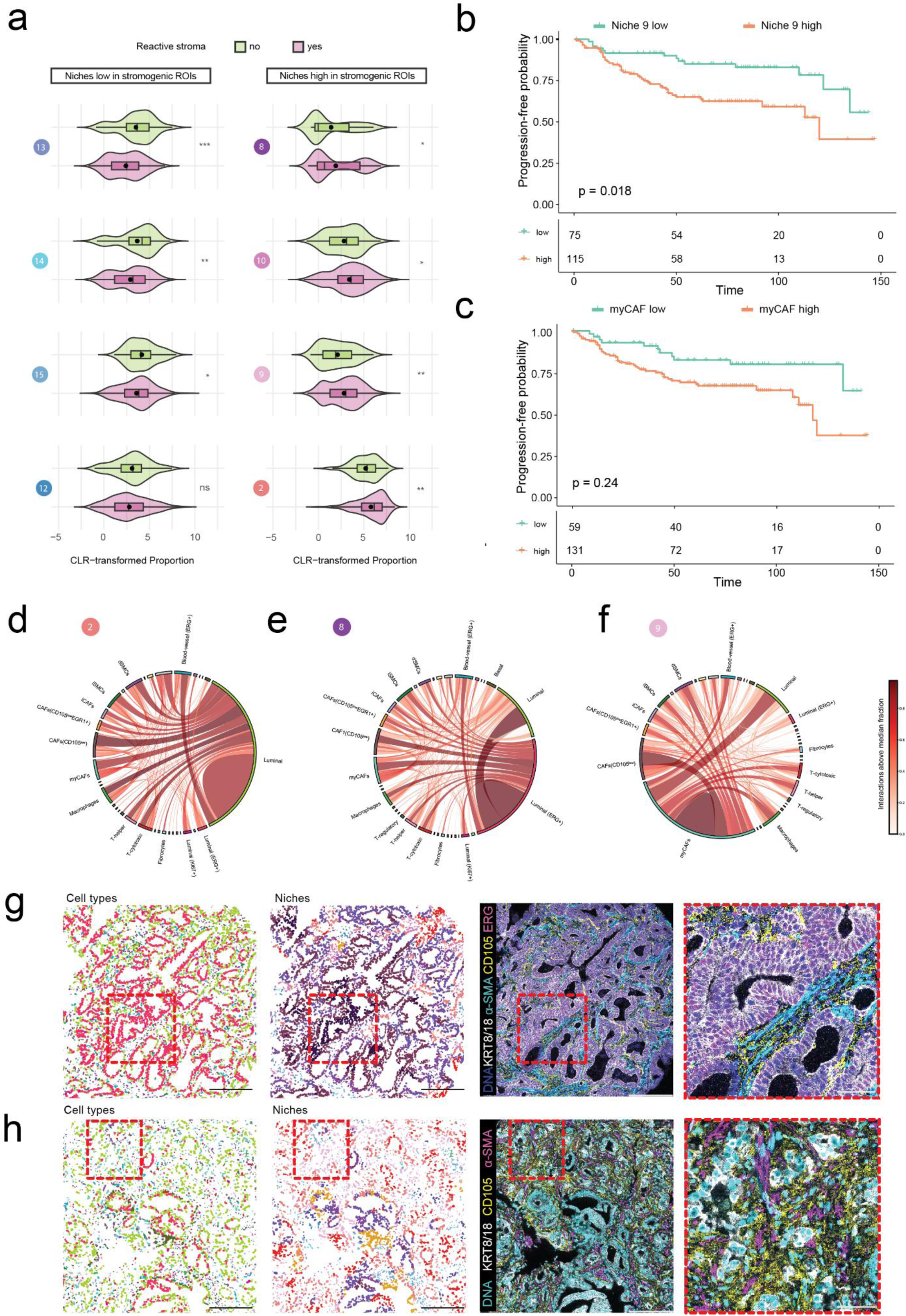
myCAF niche associates with progression-free survival and displays extensive cell-cell interactions. **a** Distribution of stromal niche abundance per stromogenic status, indicating enrichment of CAF-associated niches in stromogenic cores. Statistical significance was assessed using a Wilcoxon rank test after CLR transformation of niche proportion values, and p-values were adjusted for multiple testing using a Benjamini-Hochberg correction. **b-c** Kaplan-Meier survival analysis with patients stratified by high versus low niche 9 abundance (top) and myCAF abundance (bottom), defined in both cases at the core level using the cohort median and aggregated to the patient level (high if ≥1 high-abundance core); *p*-values were computed using a two-sided log-rank test, and adjusted for multiple testing using a Benjamini-Hochberg correction. **d-f** Circos plots of pairwise cell type interactions in niches 2, 8, 9, respectively. Edge width represents the relative abundance of interactions within each niche, and edge color indicates the consistency across cores of cell-cell interaction enrichment in the niche. **g-h** Representative cores high for niche 8 (g) and niche 9 (h), colored by cell type and niche (first two images), and IMC staining (last two images). Different features characterize these two niches in terms of cell type composition, and stroma morphology. Scale bar: 200 ∼μm (first three images). Scale bar: 50 ∼μm (magnified area, right-most images).

To explore potential mechanisms underlying the distinct behavior of these stromogenic niches, we compared cell-cell interaction patterns within niches 2, 8, and 9 (Fig. 7d-f). Niches 2 and 8 displayed similar interaction architectures, with their strongest and most consistent connections restricted to epithelial compartments (luminal epithelium in niche 2 and ERG^+^ tumor epithelium in niche 8), and relatively limited interactions with immune or other stromal subsets (Fig. 7d-e,g). Spatially, niche 8 formed a stromal rim around ERG^+^ tumor glands with limited immune infiltration (Fig. 7g). In contrast, niche 9 maintained the interaction with luminal epithelium but was further characterized by multiple high-strength interactions with immune populations (macrophages, T helper cells, fibrocytes) and other CAF subsets, including CAF CD105^low^ and SMC populations such as dSMCs (Fig. 7f, h). Together, these data suggest that the adverse prognostic impact of the myCAF stromogenic niche stems from its role as a densely interconnected stromal-immune hub, rather than from CD105 expression or stromogenic histology alone.

## Discussion

While single-cell and spatial transcriptomics have started to untangle the PCa microenvironment, its spatial organization at the protein level remains unclear^4,14,62^. Here, we present a large-scale, spatially resolved single-cell proteomic atlas of the primary PCa, integrating epithelial, stromal, endothelial, and immune phenotypes across 195 patients. We identified 18 recurrent spatial niches that reveal how localized PCa organizes into multicellular local ecosystems. Importantly, we demonstrate that spatial context provides prognostically meaningful information that cannot be inferred from cell-type abundance or Gleason grade alone.

Within the epithelial compartment, we identify an ERG^+^p53^+^ luminal cell state that robustly predicts both poor overall and progression-free survival. Strong nuclear p53 accumulation is consistent with TP53 missense mutations with adverse pathology^63^ and recent work in mouse models indicates that ERG and p53 alterations cooperate in driving aggressive disease^64^. The prognostic relevance of this co-expressing population, absent for ERG or p53 alone, supports a synergistic biological process, potentially reflecting a sequential evolutionary trajectory in which TMPRSS2-ERG fusion precedes TP53 disruption. Notably, ERG^+^p53^+^ cells were present even in low-grade regions and frequently co-occurred with immune-rich niches, suggesting that epithelial genomic instability may arise within, and be sustained by, specific inflammatory microenvironments. Recent work in mouse prostate organoids support this model, showing that ERG-overexpressing epithelial cells actively recruit and interact with macrophages, generating a ROS-rich, DNA-damage-permissive environment^65^. Additionally, mechanistic studies demonstrate that ERG reprograms double-strand break repair toward mutagenic, PARP1-dependent end-joining, and that tumor-associated macrophages foster an immunosuppressive, oxidative milieu^65–68^. Together, these observations point to a co-evolution of ERG^+^, p53-defective epithelial states with immune-enriched niches that collectively amplify genomic instability and immune evasion.

Our spatial analysis further points to substantial heterogeneity within the stromal compartment. We distinguish CAF populations with typical myCAF features, from contractile, SMC populations, and further stratify these based on CD105, CD146, CES1, EGR1 and AR expression, with the latter co-expressed in SMCs with YAP and CES1. Within the stromal compartment, CD105 (endoglin) is a notable feature: a TGFβ coreceptor enriched on angiogenic endothelium and prominently expressed by activated CAF populations. Its expression has been repeatedly linked to angiogenesis and stromal remodeling, and mechanistic studies indicate that CD105^+^ CAFs potentiate neuroendocrine differentiation and ADT resistance^39,69,70^. We have also previously shown that CD105^+^ CAFs have low AR expression and exhibit insensitivity to AR blockade, whereby disruption of the NFκB-TGFβ1-YAP1 signaling axis could curb this specific myCAF activity^39^. Importantly, findings herein demonstrate that the biological and clinical relevance of myCAF is highly dependent on spatial context: myCAFs cells preferentially organize into a small, periglandular niche enriched in histologically stromogenic regions and show the strongest association with adverse outcome. This niche exhibits extensive connectivity between myCAFs and multiple immune populations, including macrophages, T helper cells, and fibrocytes, as well as other CAF subsets, pointing to a highly interconnected stromal-immune hub coordinating multicellular interactions across compartments. Therapeutically, targeting CD105 has been proposed as a means to reduce CAF-mediated signaling and promote vascular normalization^71^. Such interventions could reprogram the stroma from a driver of immune exclusion into a microenvironment that supports effective antitumor immune responses. Our data further refine this concept by suggesting that targeting myCAFs may require consideration of their spatial organization and interacting partners, and position CD105 not simply as a marker, but as a mechanistic node at the stromal-vascular interface. As myofibroblast-like cells, CD105^+^ CAFs contribute to ECM deposition, crosslinking, and desmoplastic remodeling, leading to increased matrix stiffness, an established driver of tumor progression and poor clinical outcome in PCa^72,73^. Beyond its biomechanical effects, a stiffened ECM also acts as an immunological barrier, limiting lymphocyte infiltration and impairing anti-tumor immunity^33,74^. Immunosuppressive properties have been linked with myCAF populations implicating macrophages and perturbed T-helper cell function^75^. In this context, the adverse prognostic impact of the myCAF-rich niche (which includes direct interactions with macrophages and T-helper cells) is consistent with its role as an ECM-modifying hub that simultaneously promotes stromogenic remodeling and shapes a microenvironment hostile to adaptive immunity.

Beyond stromal organization, our analysis also identifies structured immune niches with potential functional relevance. In particular, a niche resembling a tertiary lymphoid structure (TLS) was defined by close spatial association of B cells and T cells. While canonical markers required to assess TLS maturation were not included, the observed architecture suggests the presence of organized immune aggregates. Across cancers, TLS have been associated with improved immune responses and survival, although their role in PCa appears more context-dependent^58,59,76^. Here, immune-enriched niches were preferentially associated with histologically inflamed regions, and that a composite immune niche score stratified patient outcomes, indicating that cumulative immune niche burden, rather than individual immune populations, captures prognostically relevant inflammatory states.

Taken together, our findings show that stromogenic histology alone cannot capture the clinically relevant complexity of the PCa microenvironment. Instead, spatially resolved analyses reveal that distinct multicellular architectures, ranging from the periglandular myCAF rich niche to broader epithelial, stromal, and immune ecosystems, encode prognostic information that is not evident from cellular composition alone. These results underscore the importance of integrating spatially defined CAF phenotypes and higher-order tissue organization to identify biologically meaningful tumor–stroma interfaces and improve patient risk stratification.

Several limitations should be considered. First, TLS-like structures were inferred based on spatial organization without formal assessment of maturation or localization, which may influence their clinical significance^59^. Similarly, stromogenic status was assessed using non-standardized histopathological criteria, potentially limiting its prognostic resolution^60^. Second, while IMC enables high-dimensional protein profiling, it remains restricted to a predefined marker panel and does not capture the full spectrum of cellular states, particularly within the myeloid and T cell compartments. Finally, the slow progression and relatively low number of clinical events, typical for localized PCa cohorts^77^, limits statistical power for survival analyses and underscores the need for independent external validation.

Future studies integrating IMC with spatial transcriptomics and genomic profiling will be critical to link protein-defined niches to transcriptional programs and mutational landscapes. In particular, dissecting the functional interplay between ERG⁺p53⁺ epithelial states and myCAF stromal niches may reveal actionable dependencies. Experimental models, including organoid-stroma co-culture systems, and stroma isolation^38^ will be essential to establish causal relationships and test whether genetic or pharmacologic perturbation of CD105 or downstream signaling attenuates aggressiveness and restores sensitivity to AR targeting agents.

Importantly, because IMC cannot be deployed in routine clinical diagnostics, the spatial patterns identified here may have corresponding histomorphological features detectable by more accessible assays. Linking these high-dimensional proteomic niches to conventional histopathology or targeted multiplexed approaches could enable the recognition of high-risk stromal-immune configurations without requiring full spatial proteomic profiling. This provides a potential path toward translating spatial niche information into clinically applicable biomarkers. In this context, computational approaches, including machine learning and artificial intelligence (AI) methods, may facilitate the identification of such patterns in standard H&E images, although this remains to be systematically validated.

In summary, this work reveals that spatially organized stromal-immune-epithelial ecosystems encode biologically and clinically meaningful information that is not captured by Gleason grading or bulk cell-type frequencies. The atlas generated here provides a foundation for mechanistic studies of stromal-epithelial interactions, immune modulation, and niche evolution in PCa. Integrating IMC with spatial transcriptomics, genomic profiling, and functional perturbation studies will be essential to dissect the molecular pathways underlying high-risk epithelial states and adverse stromal architectures. Beyond prostate cancer, the analytical framework established here is applicable to mapping clinically relevant spatial ecosystems across diverse tumor types.

## Materials and Methods

### Tissue Microarray Cohort

The tissue microarray (TMA) dataset includes formalin-fixed paraffin-embedded (FFPE) 1 mm^2^ prostate tissue cores obtained from 195 primary prostate cancer (PCa) patients after radical prostatectomy, provided through the European Multicenter Prostate Cancer Clinical and Translational research group (EMPaCT) study^35,36^ and the Institute of Tissue Pathology, Bern. Multiple tissue cores (up to four per patient) were taken from both high- and low-Gleason pattern prostate tissue areas to capture multifocal tissue architecture and intra-patient heterogeneity. Clinical and pathological annotations relevant for this study include patient-level Gleason grade group, biochemical recurrence (PSA progression), clinical progression (defined as local or distant recurrence), disease progression (defined as clinical progression or PSA progression), and overall survival (Supplementary Fig. 1a). Mean patient age at surgery was 66 years, with a median preoperative prostate-specific antigen (PSA) value of 37 ng/ml. Of the 195 patients included in the initial TMA cohort, five patients were removed due to the absence of tumor content across all cores. The final analytical cohort therefore comprised 190 patients with evaluable tumor regions.

### Imaging Mass Cytometry Antibody Panel Design

A 34-plex imaging mass cytometry (IMC) antibody panel was established to characterize the major compartments of the prostate TME (Fig. 1b). The panel was designed to identify: (1) epithelial cells (E-cadherin, pan-keratin, KRT8/18, KRT5, p63, androgen receptor (AR), prostate-specific antigen (PSA), synaptophysin (SYP)); (2) stromal cells including cancer-associated fibroblasts (CAFs) and smooth muscle cells (SMCs) (Vimentin, collagen type-I, *α*-smooth muscle actin (*α*-SMA), CD146, calponin-1 (CNN1), carboxylesterase 1 (CES1), CD105, early growth response-1 (EGR1)); (3) endothelial cells (CD31, CD105, podoplanin (PDPN)); (4) immune cells (CD45, CD3, CD4, CD8a, CD20, CD68, FoxP3, CD11b, CD66b); and (5) functional and pathway markers (Ki-67, cleaved caspase-3, ERG, p53, *β*-catenin, phospho-YAP1). Iridium-191/193 DNA intercalator was included to delineate nuclei for cell segmentation. CAF subpopulation markers were selected based on recent single-cell RNA sequencing (scRNA-seq) analyses of primary PCa^39,78^. CES1 was selected as an inflammatory CAF (iCAF) marker, while CD105 was selected as a marker for activated myofibroblastic CAF (myCAF) subpopulations. Phosphorylated YAP1 (pY357) was included as a marker of CAF activation state and YAP/TAZ signaling activity^39^. The stromal compartment was identified using vimentin and collagen type-I as general markers, while *α*-SMA, CD146, and CNN1 distinguished SMCs (*α*-SMA^+^, CD146^+^ CNN1^+^) from pericytes (*α*-SMA^+^, CD146^+^ CNN1^-^). Regarding immune cells, macrophages (CD68^+^), B-cells (CD20^+^), helper T-cells (CD3^+^CD4^+^), cytotoxic T-cells (CD3^+^ CD8a^+^), regulatory T-cells (CD3^+^ CD4^+^ FoxP3^+^), and polymorphonuclear myeloid-derived suppressor cells (PMN-MDSCs; CD11b^+^CD66b^+^) were selected based on their documented roles in PCa progression and ADT resistance^45,79,80^. Blood vessel endothelial cells were identified by dual CD105 and CD31 expression, while lymphatic vessels were distinguished by CD105 and PDPN co-expression. The functional markers ERG and p53 were selected for their known roles in PCa tumorigenesis^17,81,82^.

### Antibody Validation and Metal Isotope Conjugation

Carrier-free (BSA- and azide-free) antibody clones requiring in-house metal isotope conjugation were validated by immunofluorescence staining on FFPE human prostate tissue sections (cancer and normal-adjacent tissue). The antibody staining protocol matched the IMC workflow to ensure accurate performance validation. Antibodies were titrated to determine optimal dilutions. Following validation, the final IMC antibody panel was designed by selecting antibody-metal isotope pairs to minimize signal spillover among neighboring channels, considering isotope intensity and marker expression levels. Metal isotope conjugation was performed using Maxpar X8 Antibody Labeling Kits (Standard BioTools) for lanthanide metals. The procedure involved loading a specific metal isotope with the MaxPar X8 Polymer, partial antibody reduction with Tris(2-carboxyethyl)phosphine hydrochloride (TCEP), and conjugation of the partially-reduced antibody with the lanthanide-loaded polymer. Conjugated antibody concentrations were measured using a NanoDrop 2000/2000c Spectrophotometer. Metal-tagged antibodies were eluted at 0.5 mg/ml in Antibody Stabilizer PBS (Boca Scientific) supplemented with 0.05% sodium azide and stored at 4°C.

### Imaging Mass Cytometry Staining

IMC antibody staining was performed with preconjugated metal-tagged antibodies (Standard BioTools) or conjugated in-house. All antibodies used are described in Supplementary Table 1. The IMC antibody panel was first titrated on human PCa tissue samples to select optimal dilutions. Freshly cut FFPE tissue sections (not older than one day) from TMA blocks were used. TMA blocks were sectioned at the Translational Research Unit (TRU) at the Institute of Tissue Medicine and Pathology, University of Bern. Tissue slides were incubated for 2 hours at 60°C to remove visible wax and enhance tissue adhesion. Deparaffinization was performed by sequential transfers through xylol, decreasing ethanol concentrations (100%, 95%, 80%, and 70%), and distilled H_2_O. Denatured ethanol (90% absolute ethanol + 5% isopropanol + 5% methanol) was used. Antigen retrieval was performed with Antigen Retrieval Buffer (Agilent) at pH 9 for 30 min. Blocking was performed in 3% bovine serum albumin (BSA) dissolved in PBS 1x. Metal-conjugated antibodies were incubated overnight at 4°C in 0.5% BSA/PBS solution. The Maxpar IMC Cell Segmentation Kit, containing three individual plasma membrane markers conjugated to metal isotopes (ICSK1, ICSK2, ICSK3), was added to the antibody mix. Washing steps were performed with 0.2% Triton X-100 in PBS and PBS solutions. Iridium (191Ir, 193Ir) DNA intercalating agents (Cell-ID Intercalator-Ir, Standard BioTools) were incubated for 30 minutes at room temperature in PBS 1x. Final washes were performed in PBS 1x and distilled H_2_O. Stained slides were air-dried and stored in a dry environment until IMC acquisition (slides can be stored for up to 1 year). MaxPar PBS and distilled MaxPar Water (Standard BioTools) were used throughout the IMC staining protocol.

### H&E Staining

H&E staining was performed on TMA FFPE serial sections adjacent to those used for IMC staining to enable histopathological assessment. Following deparaffinization through xylol and decreasing ethanol concentrations, tissues were incubated with hematoxylin (marking cell nuclei) and eosin (marking basic cytoplasmic and extracellular matrix proteins). Tissues were then processed through increasing ethanol concentrations and xylol, and mounted with Entellan/Eukitt Mounting medium (Sigma Aldrich). Images were acquired using the Panoramic 250 Flash II Slide Scanner (3DHistech). An independent pathological evaluation of H&E-stained sections provided core-level annotations, including Gleason group, presence of stromogenic reaction, and inflammation status.

### IMC Data Acquisition

IMC data were acquired at the Imaging Mass Cytometry and Mass Cytometry Platform of the University of Bern and Inselspital, Bern. Stained and dried samples were inserted into the Hyperion Tissue Imager (Standard BioTools), where a 1 μm diameter UV laser ablated the tissue within selected regions of interest (ROIs) at a raster rate of 200 Hz. With each laser shot, the tissue at the ablation spot was vaporized, and the resulting plume containing heavy metals from the tissue was transported into the Helios mass cytometer. Elements were atomized and ionized in the inductively coupled plasma, then pulsed into the time-of-flight (TOF) chamber for detection based on their mass-to-charge ratio. Mass cytometry data (MCD) files were acquired using CyTOF software version 7.0.8493. The average tissue area of each ROI was approximately 1 mm^2^ (area of a TMA core). A total of 523 ROIs were acquired across the cohort. ROIs lacking clinical annotations or tumor content were removed, resulting in a final dataset of 459 tumor-containing ROIs.

### Spillover Compensation

For spillover assessment, a 2% agarose solution in distilled H_2_O was prepared, and microscope glass slides were dipped in the agarose solution and allowed to dry completely (at least for 30 minutes). Arrays of 0.3 μl drops of 0.4% trypan blue dye were spotted onto the agarose-coated slide, and 0.3 μl of each specific metal-tagged antibody was added to a unique trypan blue spot. The slide was dried for 1 hour and then acquired with the Hyperion Tissue Imager (Standard BioTools) to assess channel spillover.

### Data Processing

Raw IMC data underwent minimal processing using Steinbock (version 0.16.3) to remove hot pixels^83^. Channel spillover was evaluated with the Bioconductor package CATALYST (version 1.26.1)^84^, which indicated negligible crosstalk requiring no further corrections. Due to non-specific staining following in-house metal isotope conjugation, the fibroblast activation protein (FAP) marker was excluded from subsequent analyses. Images underwent nuclear segmentation using Steinbock and DeepCell^85^ based on the DNA1 and DNA2 channels (Iridium 191/193). Nuclear segmentation was employed due to challenges in defining cellular borders for CAFs and SMCs, which display elongated morphology and lack specific membrane protein markers. Nonetheless, extraction of protein expression data from each nuclear mask was sufficient to distinguish major cell types and their subclusters. Segmentation results were visually inspected, and regions presenting staining artifacts or obvious segmentation errors were removed. This approach yielded a total of 2’191’967 single-cell profiles across all ROIs. Single-cell marker expression was determined by calculating the mean pixel intensity within each segmentation mask. For all downstream data analysis and visualizations, per-cell marker values were clipped at the 99.9^th^ percentile across the entire dataset and underwent arcsinh transformation (cofactor = 1). The arcsinh-transformed single-cell values were then min-max normalized to a [0-1] range.

### Cell Phenotyping

Cell-type annotation was performed using an iterative clustering and marker-based annotation strategy. First, all cells were embedded and clustered in an unsupervised manner: a k-nearest neighbors (k-NN) graph of all cells was computed using the rapids_singlecell (version 0.4.0) implementation of Parametric UMAP^86^, followed by community detection using the Leiden algorithm^87^, as implemented in the igraph package (https://igraph.org).

Clusters were subsequently annotated based on canonical marker expression profiles. To increase phenotypic resolution, we applied a hierarchical refinement strategy in which major cellular compartments were first identified and then re-clustered independently. At the top level, clusters were grouped into immune and non-immune populations based on the expression of canonical immune markers (e.g., CD3, CD4, CD8a, CD20, CD68, FoxP3, CD11b, CD66b). The immune compartment was then re-clustered using all immune-specific markers, yielding 11 distinct immune cell populations (Fig. 2a).

The non-immune compartment was further subdivided into epithelial, endothelial, and stromal lineages based on established markers. Epithelial cells were identified using pan-keratin, E-cadherin, KRT8/18, KRT5, and p63, and further resolved into nine phenotypic groups using lineage and functional markers (e.g., AR, PSA, Ki-67, p53, YAP1). A focused re-clustering of basal-like populations improved separation into basal, luminal, and transitional states. Endothelial cells were defined by CD31 and ERG expression and further subdivided into three phenotypes. Remaining non-immune, non-endothelial cells were assigned to the stromal compartment and re-clustered using mesenchymal and fibroblast-associated markers (e.g., α-SMA, collagen I, vimentin, CD146, CD105, PDPN), while excluding CD45+ and CD31+ populations. This approach identified multiple stromal subpopulations, forming the basis for downstream analysis of fibroblast heterogeneity.

Cells that could not be confidently assigned at intermediate stages were labeled as undefined and re-clustered using the full marker panel. Due to the small size of immune cells and the imaging resolution (∼1 µm), a minority of cells exhibited apparent double positivity (e.g., CD4⁺CD20⁺), likely reflecting segmentation limitations in densely packed regions rather than true co-expression.

Overall, this iterative framework combines unsupervised clustering with marker-based annotation to achieve robust and biologically consistent cell-type classification while preserving phenotypic granularity. Hierarchical clustering of normalized mean marker expression across the 34 annotated cell types (Fig. 2a) was performed using a Euclidean distance metric and average linkage.

### Cell Type Proportion Quantification

Cell type proportions were computed per core after adding a pseudocount of 1 to all cell type to avoid zero values in downstream log-ratio transformations. To identify groups of patients with similar cellular compositions (Fig. 4a) we derived a patient-level composition vector by aggregating core-level proportions by max-pooling (*i.e.*, selecting the maximum observed proportion of each cell type across all cores from the same patient). Patient-level compositions were then clustered by hierarchical clustering using average linkage and a Jensen-Shannon divergence as a distance metric between patient compositions. The same clustering process but without aggregation was followed for the core-level groups (Supplementary Fig. 3).

To enable downstream regression modeling, compositional data were transformed from the simplex to an unconstrained real space using the centered log-ratio (CLR) transformation:

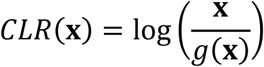

where 𝐱 is the vector of cell type proportions and 𝑔(𝐱) is the geometric mean of all proportions. This transformation addresses the unit-sum constraint inherent in compositional data and creates log-ratio coordinates appropriate for standard statistical methods. The CLR-transformed values were used as continuous predictors in all subsequent survival analyses.

### Cox Proportional Hazards Regression

Univariate Cox proportional hazards regression was performed to assess associations between CLR-transformed cell type proportions and overall survival or disease progression (Fig. 4d). Survival time was defined as the interval from diagnosis to the event of interest (death or disease progression) or last follow-up. For each cell type, a separate Cox model was fit with the CLR-transformed proportion as the sole predictor:

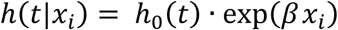

where 𝑥_𝑖_denotes the CLR-transformed proportion of cell type 𝑖 , ℎ(𝑡|𝑥_𝑖_) is the hazard function at time 𝑡, ℎ_0_(𝑡) is the baseline hazard, and 𝛽 is the log hazard ratio associated with a one-unit increase in 𝑥_𝑖_. Hazard ratios (HR) with 95% confidence intervals were computed for each model. To account for multiple testing across cell types, false discovery rate (FDR) correction was applied using the Benjamini-Hochberg procedure. Statistical significance was defined as FDR-adjusted 𝑝 < 0.05. All Cox regression analyses were conducted using the survival package in R.

### Spatial Niche Identification

Tissue niches were identified using a neighborhood-based approach adapted from previous study^57^. For each IMC core, a cell-cell proximity graph was generated: each cell 𝑖 was connected to all neighboring cells within a 32-pixel radius (≈32 𝜇m). The local neighborhood composition of cell 𝑖 was quantified as the vector 𝑓_𝑖_ = (𝑓_𝑖1_, 𝑓_𝑖2_, … , 𝑓_𝑖𝑇_), where *T* denotes the number of cell types and 𝑓_𝑖𝑡_ is the frequency of cell type 𝑡 in the local neighborhood of cell 𝑖. Cells with fewer than 3 interactions were discarded from the analysis. This procedure resulted in a 𝑛_𝑐𝑒𝑙𝑙𝑠_ × 𝑇 matrix 𝐹 that captures local neighborhood representations across the PCa TME. These local representations were subsequently clustered to identify recurrent multi-cell interaction patterns corresponding to putative TME niches. Clustering was performed using k-means (scikit-learn implementation) with a Euclidean distance metric and k-means++ initialization, testing k = 24 clusters. To avoid spurious niche definitions, a post-clustering filtering step retained only clusters present in at least 5 patients and containing at least 15 cells per sample occurrence.

To evaluate clustering robustness, k-means was repeated 50 times with different random seeds, and clustering partitions were compared using the adjusted Rand index (ARI)^88^, as implemented in scikit-learn. The ARI quantifies the probability that a pair of cells is assigned to the same niche in both runs, adjusting for chance. An ARI of 1 indicates identical niche assignments, whereas values close to zero indicate random assignments. The identified niches showed high agreement between runs (Supplementary Fig.7a). The final niche assignment was selected as the run with maximal overall agreement to all other runs (highlighted in Supplementary Fig.7a). One niche was excluded after filtering, and the remaining 23 niches were manually validated based on cell-type composition and spatial co-localization, and further examined for biological plausibility, leading to the merging of five niches, and yielding a final set of 18 distinct niches.

### Niche analysis and cell-cell interactions

Cell-type enrichment within spatial niches was quantified using z-scores. For each cell type, the mean frequency within a given niche was compared to its mean frequency across all samples and normalized by the corresponding standard deviation across samples. This metric captures relative over- or under-representation of cell types within each niche compared to their baseline abundance. Niche-level cell type enrichment profiles were then clustered using Spearman correlation as a distance metric to group niches with similar cellular compositions into higher-order niche categories (Fig. 5a).

Niche co-occurrence within tumor cores was quantified by computing pairwise Spearman correlation coefficients between niche abundance scores across cores. Positive and negative correlations indicate co-occurrence and mutual exclusion, respectively. The resulting correlation matrix was clustered using Euclidean distance to identify higher-order patterns of niche co-occurrence (Fig. 5b). Patient clustering per niche abundance (Fig. 6a) was done using a Jensen-Shannon Divergence (JSD) distance between patient-level niche compositions.

Cell-cell interactions were defined based on radius graph connectivity. For each niche, pairwise interaction frequencies between cell types were first quantified on a per-core basis and normalized by the total number of interactions within that niche, considering only cores with ≥15 niche cells. Interaction frequencies were then averaged across cores in which the niche was present. To assess the consistency of interactions across cores, niche-specific interaction frequencies were compared to a cohort-wide median interaction frequency for each interaction. The cohort-wide median was computed from core-wise normalized interaction frequencies aggregated across all cores irrespective of niche annotation. For each pairwise interaction, the proportion of cores in which the interaction frequency within the niche exceeded this median was calculated. These two metrics were used to construct circos plots (Fig. 7d-f), where edge width represents normalized interaction frequency within the niche, and edge color reflects the proportion of cores with overrepresentation of the interaction within the niche.

### Kaplan-Meier Survival Analysis

Kaplan-Meier survival curves were used to assess associations between clinical outcomes (overall survival and progression-free survival) and features derived from cell-type composition, spatial niche organization, composite niche scores, and histopathological annotations. Survival time was defined as the interval from diagnosis to event (death or disease progression) or last follow-up. For each feature of interest, patients were stratified into groups based on predefined criteria.

For features quantified at the core level (e.g., cell-type or niche abundance), patients were stratified based on aggregated core-level measurements. Specifically, each core was classified as high or low for the cell type or niche of interest based on the cohort-wide median proportion of the respective cell type/niche, computed across cores with non-zero cell type/niche abundance. Patient-level groups were then defined by aggregating core-level labels: a patient was assigned to the high group if at least one core was classified as high, and to the low group otherwise.

For categorical core-level annotations, namely inflammation and stromogenic status, patients were stratified as positive or negative based on the presence of the respective positive label in at least one patient core.

For composite metrics, such as the immune niche-based risk score (Fig. 6e), patients were stratified based on the number of immune-associated niches (niches 16-18) present at high relative abundance. For each niche, high abundance was defined as exceeding the 75^th^ percentile of its distribution across all cores in the cohort, computed across cores with non-zero abundance of the respective niche. Patients were then assigned to one of four groups (0, 1, 2, or 3) according to how many of these niches (0, 1, 2, or all 3) reached high abundance in at least one core.

For all Kaplan-Meier analyses, survival probabilities were estimated at each observed event time, and survival curves were compared using a two-sided log-rank test. Confidence intervals for survival probabilities were computed using Greenwood’s formula. Risk tables showing the number of patients at risk at each time point were included. All Kaplan-Meier analyses were performed using the survival and survminer packages in R.

### Statistical Software

All statistical analyses and data visualizations were performed in R (version 4.4.2) and Python (versions 3.12.12 and 3.11.11). Key packages used include: survival and survminer for survival analysis; seaborn, matplotlib, ggplot2, umap-learn, ComplexHeatmap for visualization; composition and base R packages for statistical analyses, athena, scikit-learn, rapids_singlecell, and igraph for clustering and graph/spatial analyses; and Steinbock for IMC data preprocessing.

## Acknowledgements

This project was funded by Swiss National Science Foundation (SNSF) grants #189369, #202297, #189149 to MK, #202297 to MR and an Austrian Science Fund (FWF) grant doi: 10.55776/I4565 to NS. The authors would like to acknowledge the Imaging Mass Cytometry Platform of the University of Bern and Inselspital (Bern), as well as the Translational Research Unit (TRU) at the Institute of Tissue Medicine and Pathology (University of Bern). Also, the clinical study coordinator of the Insel Urology Department in Bern Ms Anselm Lafita for the excellent work.

## Author contributions

M.R. and M.K.D.J. conceived and designed the study. F.B. and S.K. developed the imaging mass cytometry (IMC) antibody panel, and conducted IMC experiments. E.B., E.D, L.N. and N.S. contributed to the IMC panel design and performed marker validation experiments. M.S. provided clinical resources and cohort access. E.C. performed histopathological evaluation and clinical annotation of tissue samples. A.M. and E.M. developed the computational pipeline, and performed data analysis. A.M., A.B., F.B., M.E., S.K., M.R., M.K.D.J. interpreted the data, with contributions from G.N.T., A.L. and B.R. A.B., A.M. and M.E. generated and prepared manuscript figures. A.B., M.R., M.K.D.J, and S.K. wrote the manuscript with input from all authors. M.R. and M.K.D.J. supervised the study. All authors reviewed and approved the final manuscript.

## Data availability

The imaging mass cytometry (IMC) data generated in this study, including single-cell expression matrices, cell-type annotations, and spatial niche assignments, have been deposited in Zenodo (10.5281/zenodo.19552665) for academic non-commercial research. Patient clinical response data will be made available upon request to the corresponding authors.

## Code availability

All custom code used for data processing, single-cell analysis, spatial niche identification, and statistical analysis is publicly available at GitHub: https://github.com/AI4SCR/prostate-cancer-microenvironment. All scripts necessary to reproduce the main figures and results are provided in the repository.

---

**Supplementary Figure 1.**
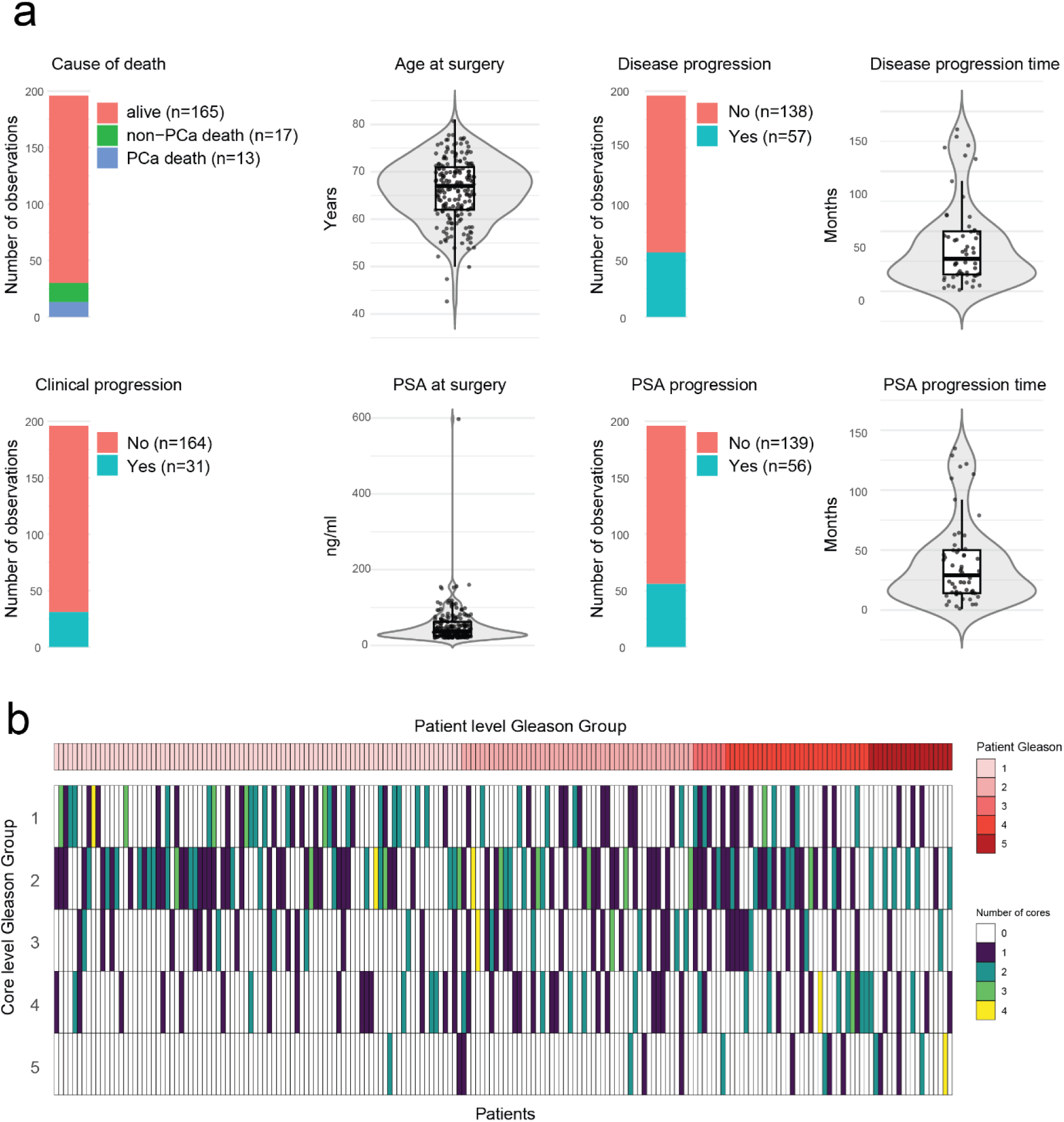
EMPaCT cohort description. **a** Cohort metadata summary, showing the distribution of the available patient metadata (cause of death, age at surgery, PSA at surgery) and clinical follow-up variables (PSA-based and/or clinical disease progression with time-to-event in months). **b** Core-to-patient Gleason group concordance heatmap, ordered by increasing patient-level Gleason group (top), where each column represents the Gleason group distribution per patient across all TMA cores (up to 4 cores per patient).

**Supplementary Figure 2:**
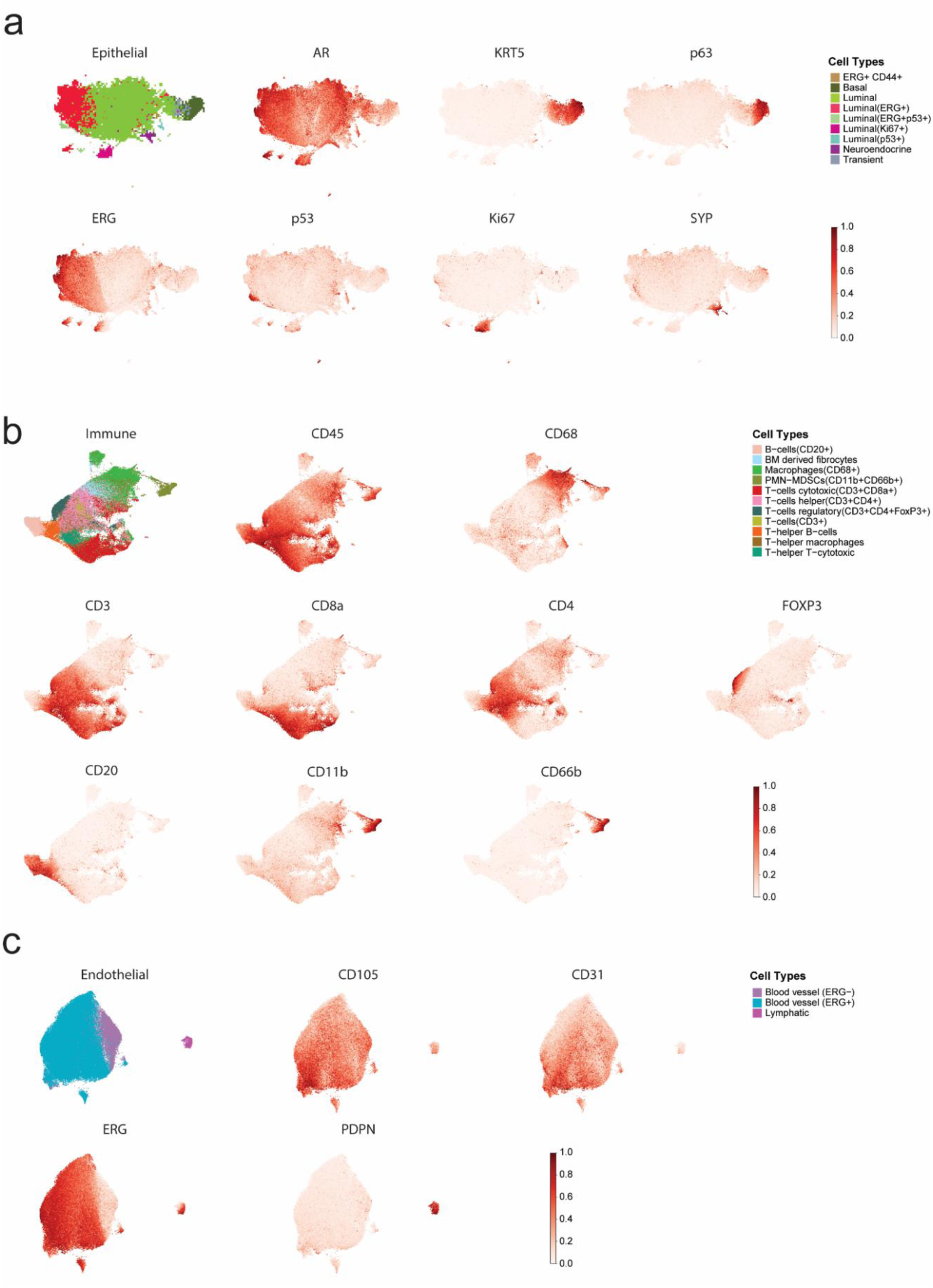
Marker expression in compartment-specific UMAP embeddings. **a-c** UMAPs of epithelial, immune, and endothelial compartments, respectively, colored by cell type (left) and selected marker intensities.

**Supplementary Figure 3:**
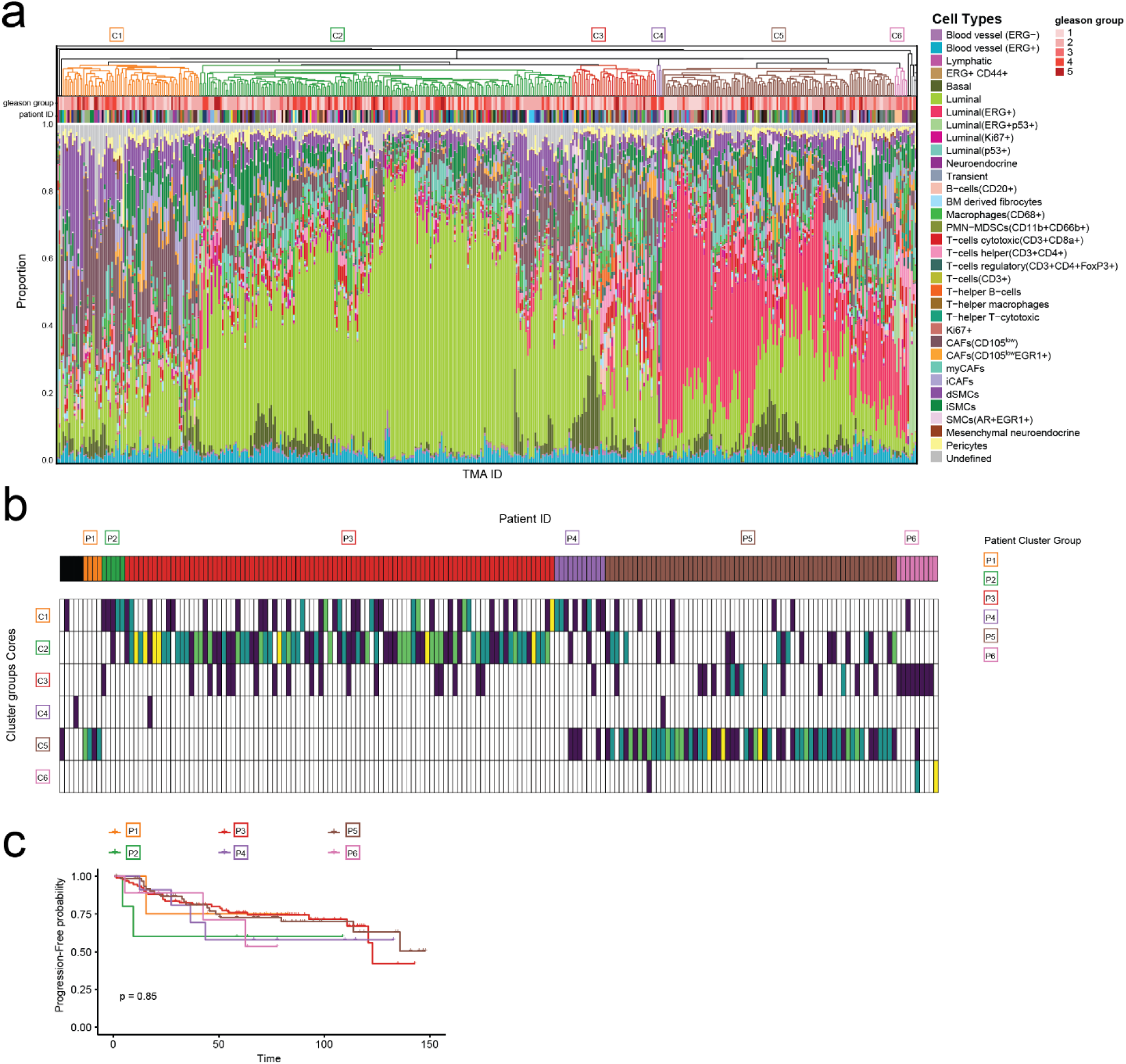
Core-level clustering of cell-type compositions. **a** Hierarchical clustering of core-level cell-type proportions, demonstrating six core-level groups. **b** Core-to-patient cell composition clustering concordance heatmap, ordered by patient-level clusters (top), where each column represents the core-level cluster distribution per patient across all TMA cores (up to 4 cores per patient). **c** Kaplan-Meier analysis of progression-free survival probability stratified by patient group; statistical significance was assessed using a two-sided log-rank test.

**Supplementary Figure 4:**
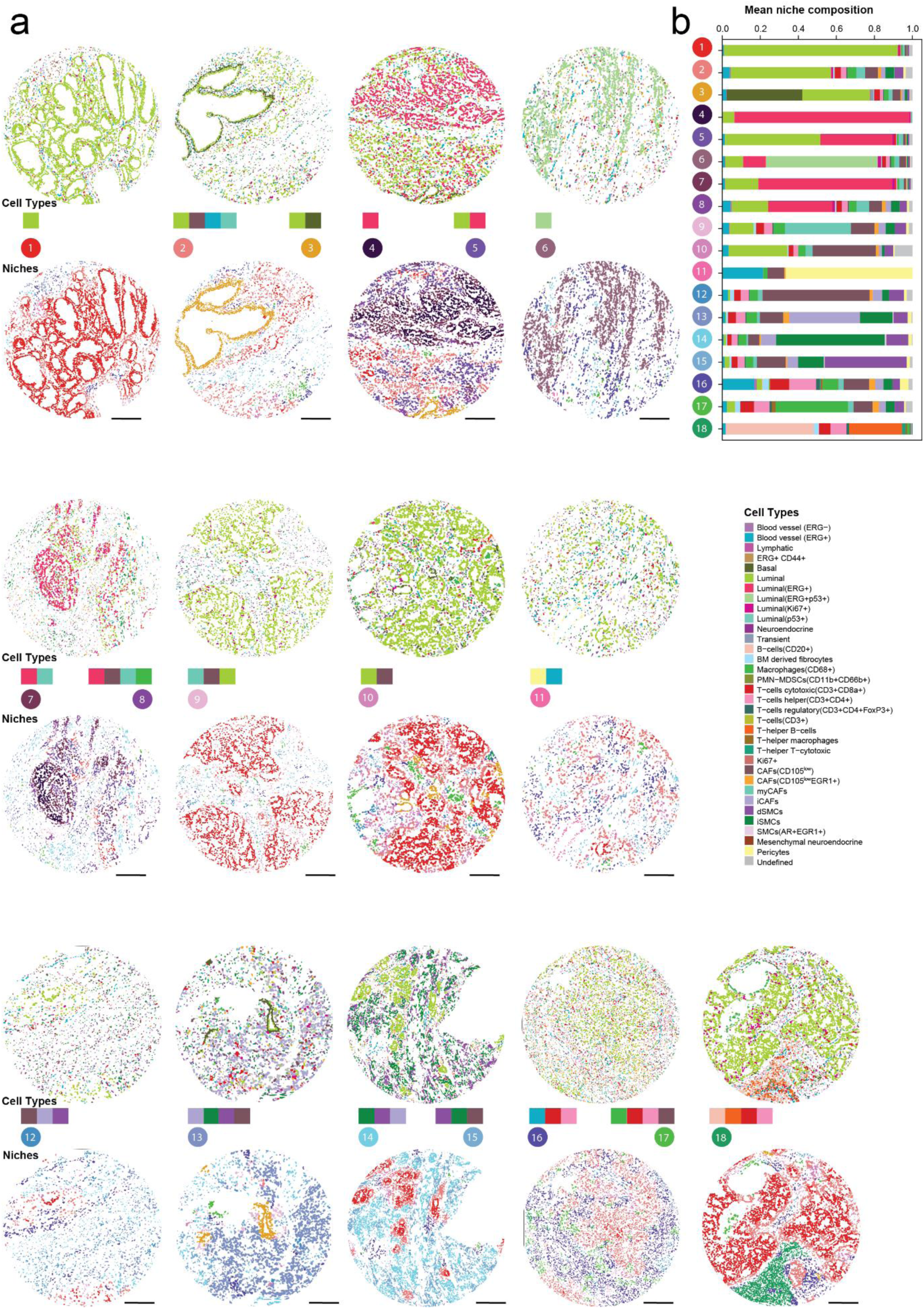
Niche composition and spatial organization. **a** Representative images of all niches, with matching cell type composition on matching ROI images. Scale bar: 200 ∼μm. **b** Stacked barplot of cell type composition of the identified niches.

**Supplementary Figure 5.**
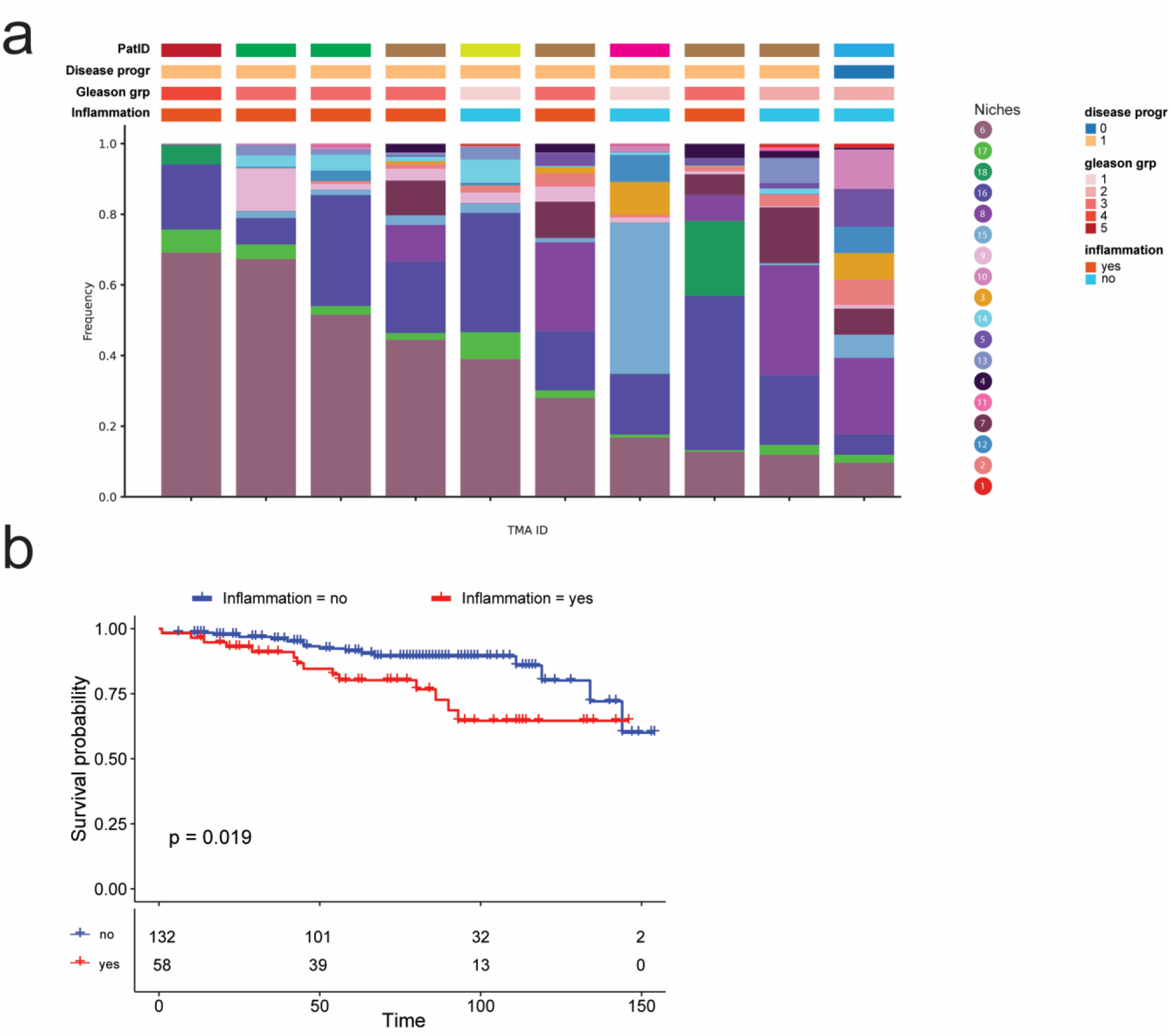
Inflammatory niches co-occur with ERG^+^P53^+^ tumor niche and associate with worse survival. **a** Stacked barplot of cell type composition of niche 6 ordered by its proportion in each core. **b** Kaplan-Meier survival analysis with patients stratified by inflammation status, defined by the presence of at least one core annotated as inflamed; *p*-values were computed using a two-sided log-rank test.

**Supplementary Figure 6.**
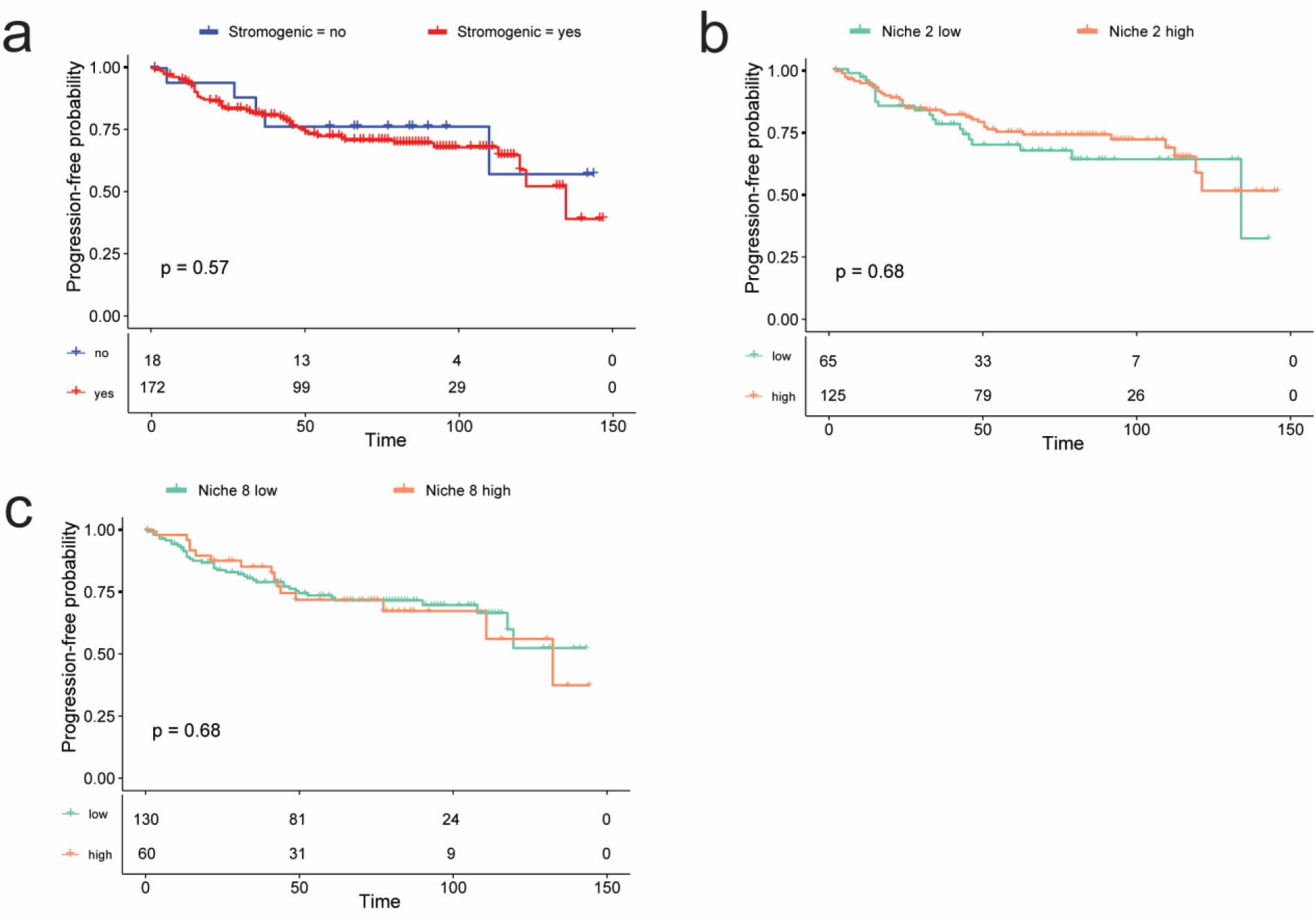
Niche 9 associates with worse clinical outcomes. **a** Kaplan-Meier progression-free survival analysis with patients stratified by stromogenic status, defined by the presence of at least one core annotated as stromogenic; *p*-values were computed using a two-sided log-rank test. **b-c** Kaplan-Meier progression-free survival analysis for abundance of niche 2 and 8, respectively. Patients were grouped into high and low categories based on niche abundance, defined at the core level using the cohort median and aggregated to the patient level (high if ≥1 high-abundance core); p-values were computed using a two-sided log-rank test.

**Supplementary Figure 7.**
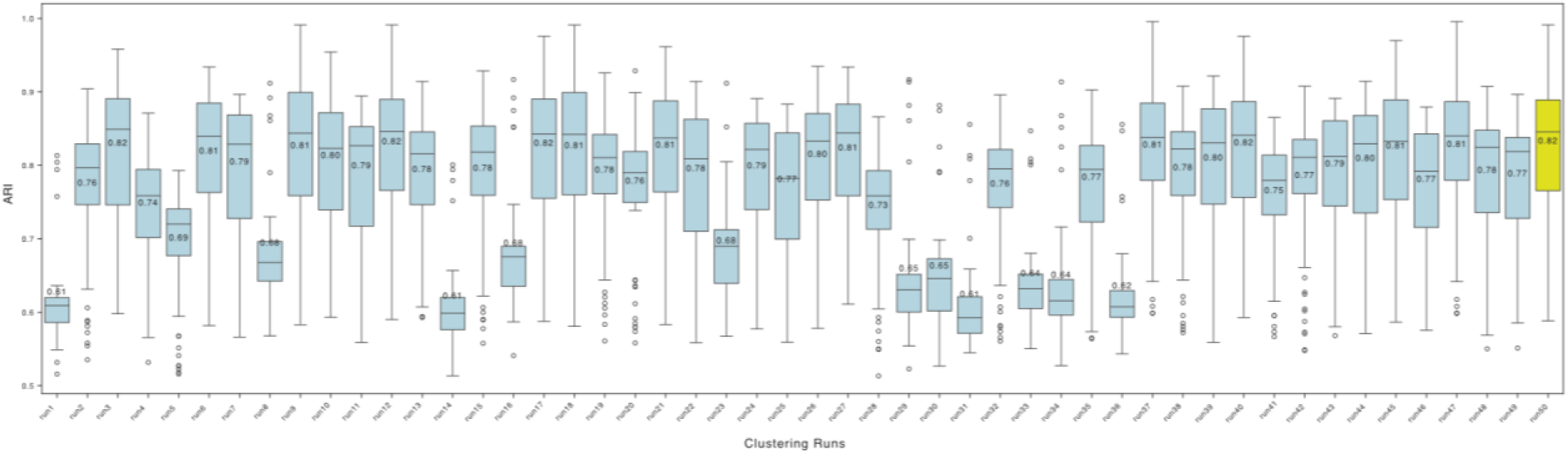
Robustness of tissue niche clustering assessed by adjusted Rand index (ARI). K-means clustering was repeated 50 times using different random seeds. For each run, clustering agreement with all other runs was quantified using the adjusted Rand index (ARI). Each boxplot shows the distribution of pairwise ARI values between one clustering solution and all other runs. Higher ARI values indicate greater similarity between clustering solutions. The clustering run with the highest overall agreement across all comparisons is highlighted in yellow and was selected as the final solution for downstream analyses.

**Supplementary Table 1.**
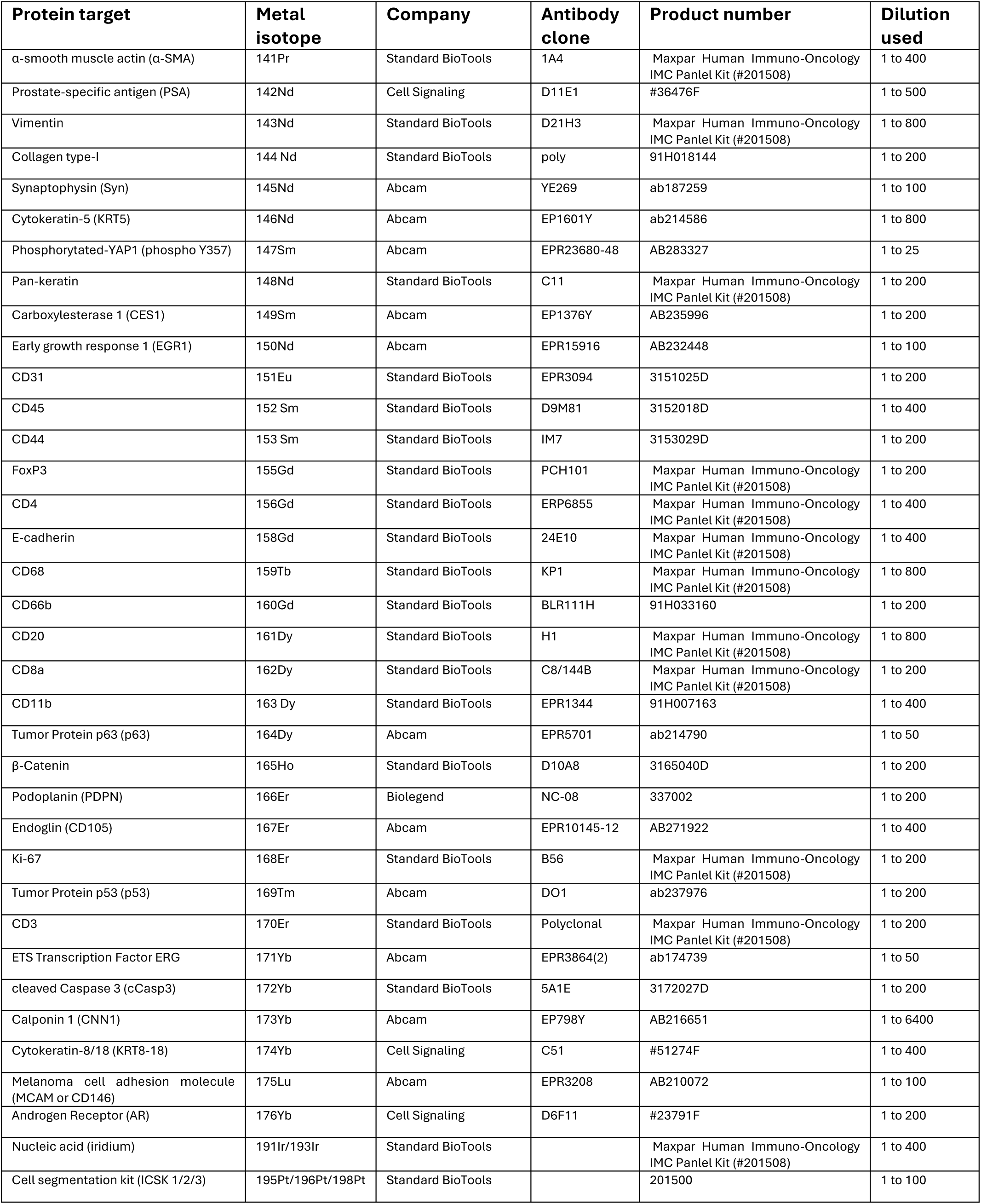
Description of the IMC antibody panel used.

