## Supplementary Figures for "Spatial single-cell proteomics defines multicellular niches in the primary prostate cancer microenvironment"

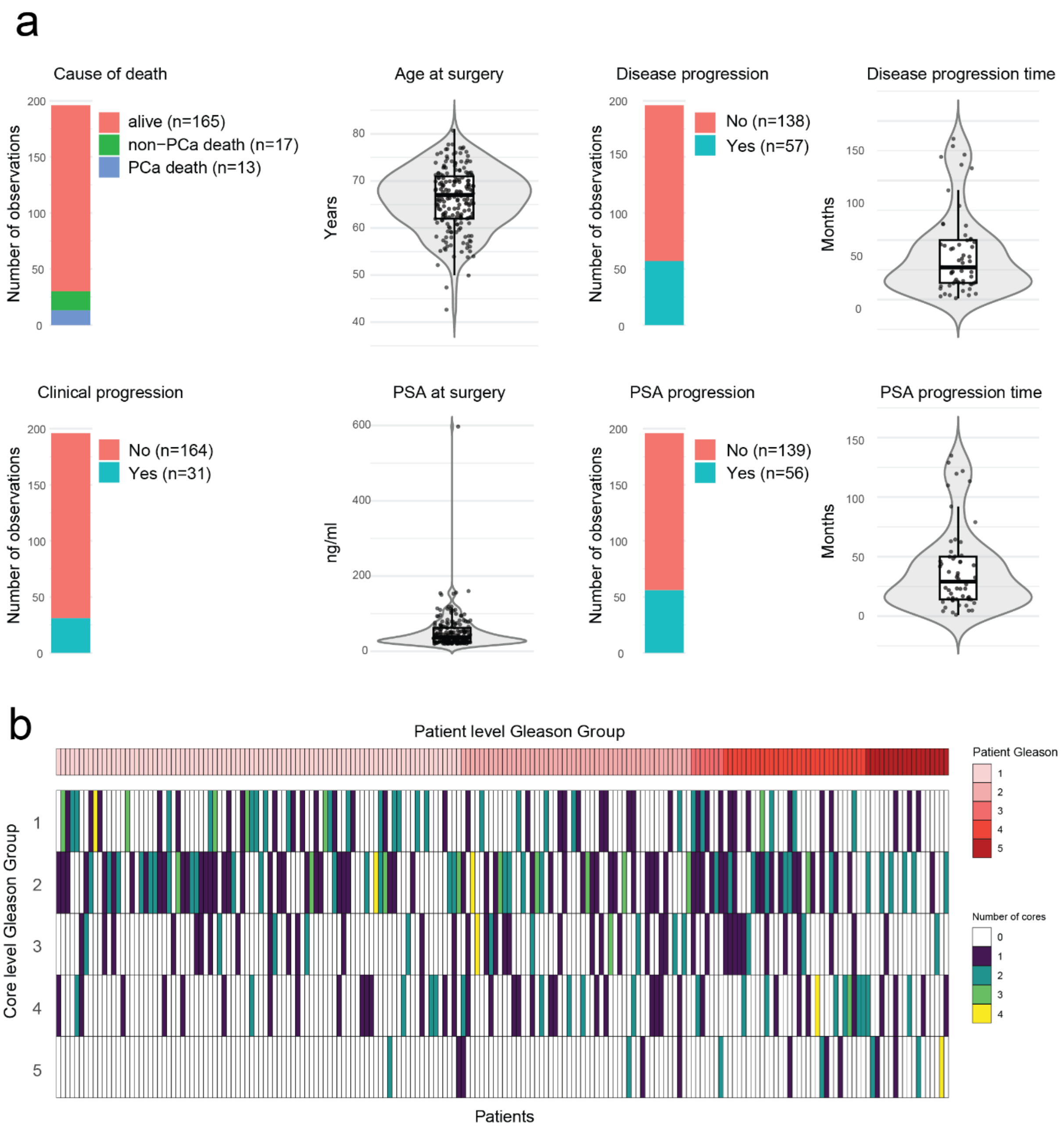


**Supplementary Figure 1. EMPaCT cohort description**. **a** Cohort metadata summary, showing the distribution of the available patient metadata (cause of death, age at surgery, PSA at surgery) and clinical follow-up variables (PSA-based and/or clinical disease progression with time-to-event in months). **b** Core-to-patient Gleason group concordance heatmap, ordered by increasing patient-level Gleason group (top), where each column represents the Gleason group distribution per patient across all TMA cores (up to 4 cores per patient).

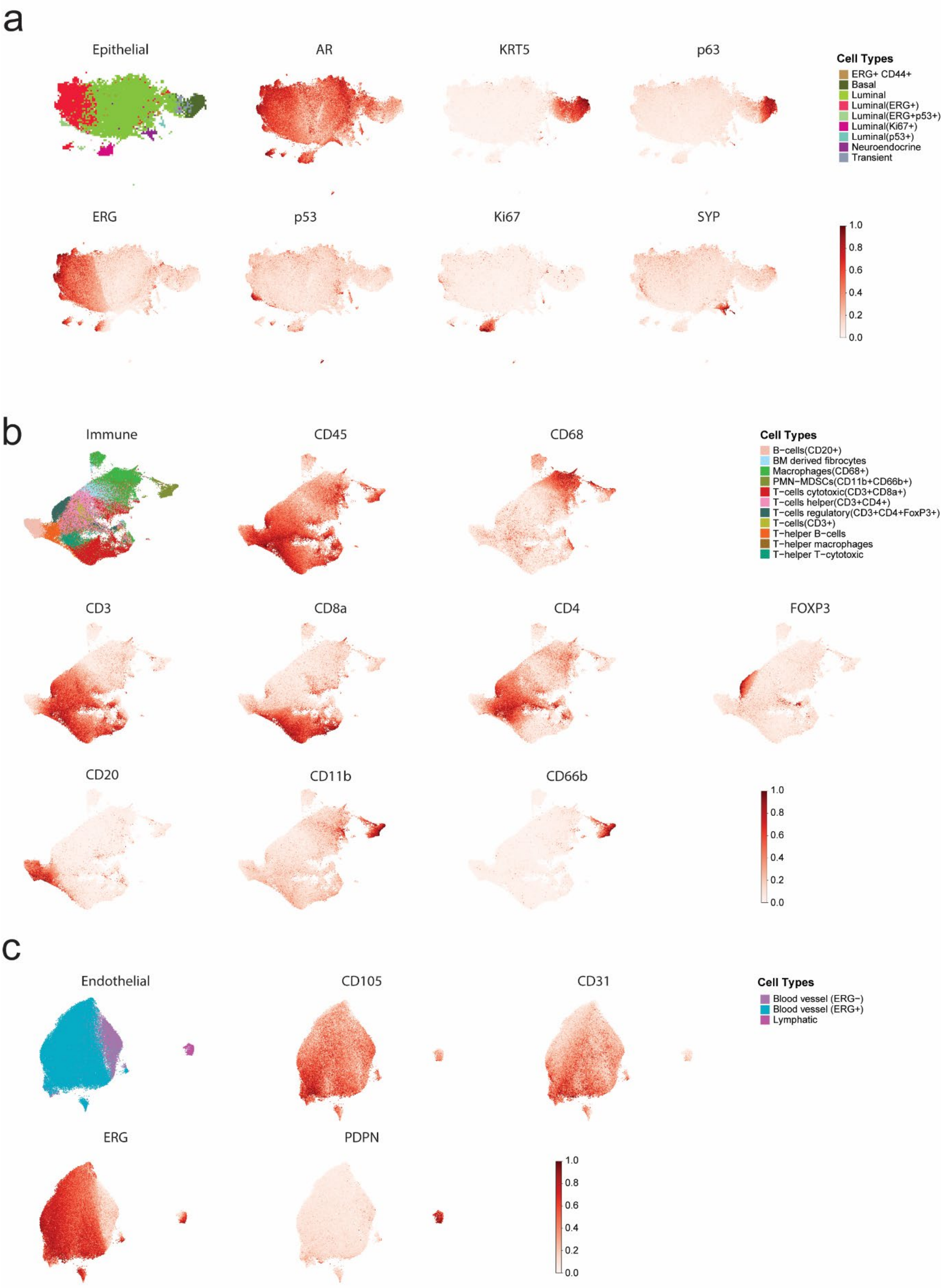


**Supplementary Figure 2: Marker expression in compartment-specific UMAP embeddings. a-c** UMAPs of epithelial, immune, and endothelial compartments, respectively, colored by cell type (left) and selected marker intensities.

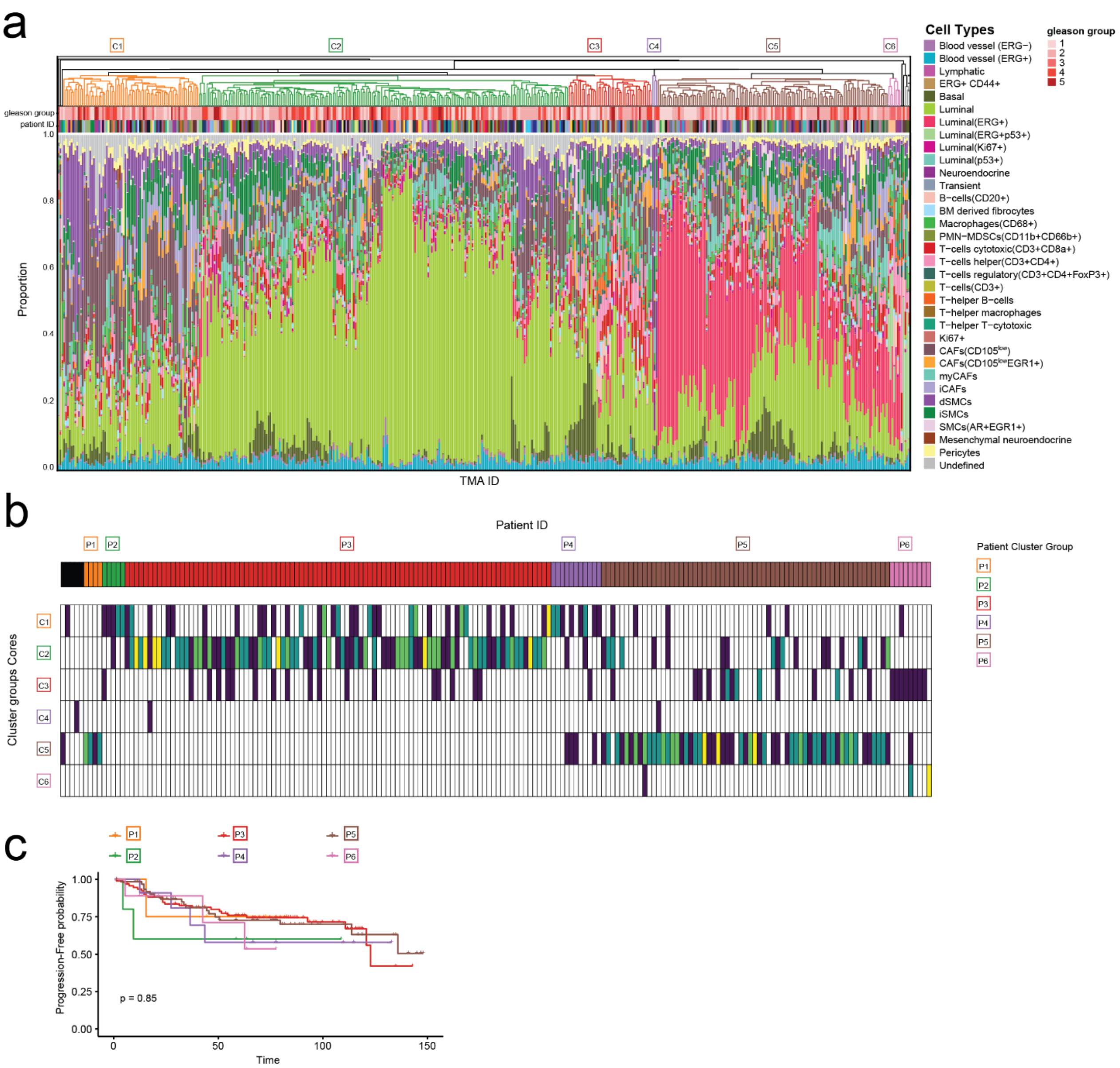


**Supplementary Figure 3: Core-level clustering of cell-type compositions. a** Hierarchical clustering of core-level cell-type proportions, demonstrating six core-level groups. **b** Core-to-patient cell composition clustering concordance heatmap, ordered by patient-level clusters (top), where each column represents the core-level cluster distribution per patient across all TMA cores (up to 4 cores per patient). **c** Kaplan-Meier analysis of progression-free survival probability stratified by patient group; statistical significance was assessed using a two-sided log-rank test.

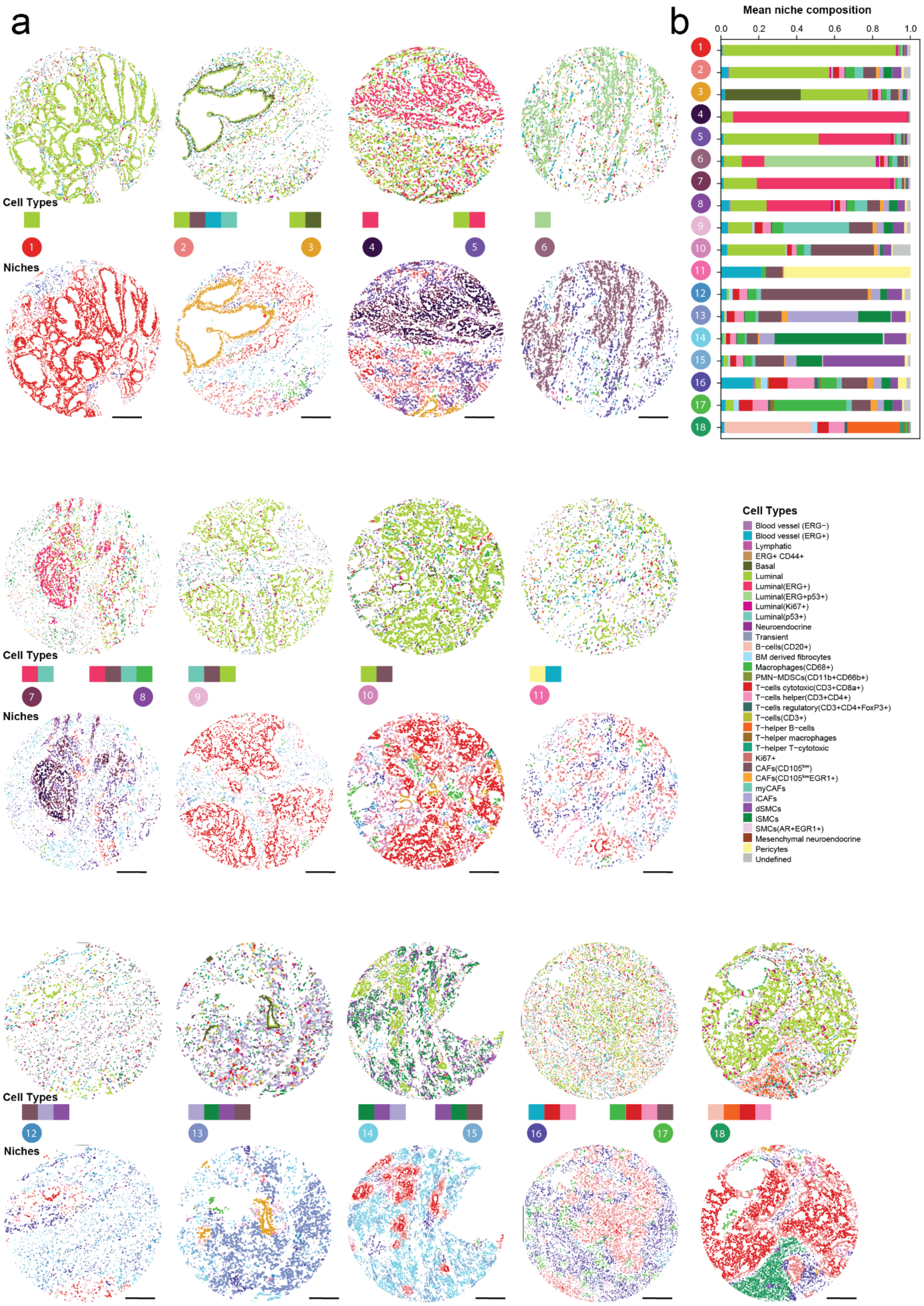


**Supplementary Figure 4**: **Niche composition and spatial organization.** **a** Representative images of all niches, with matching cell type composition on matching ROI images. Scale bar: 200 ~µm. **b** Stacked barplot of cell type composition of the identified niches.

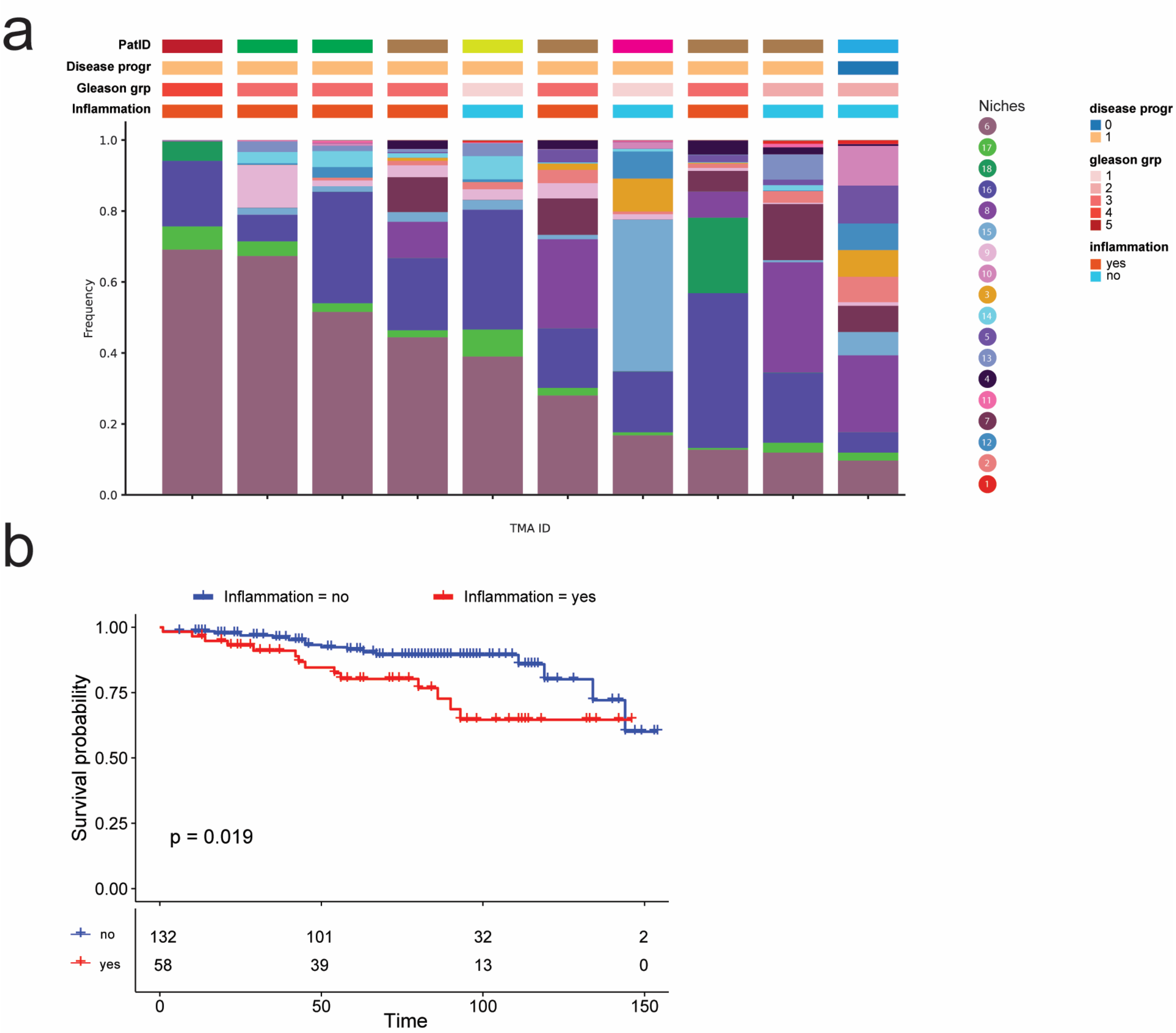


**Supplementary Figure 5. Inflammatory niches co-occur with ERG⁺P53⁺ tumor niche and associate with worse survival. a** Stacked barplot of cell type composition of niche 6 ordered by its proportion in each core. **b** Kaplan-Meier survival analysis with patients stratified by inflammation status, defined by the presence of at least one core annotated as inflamed; *p*-values were computed using a two-sided log-rank test.

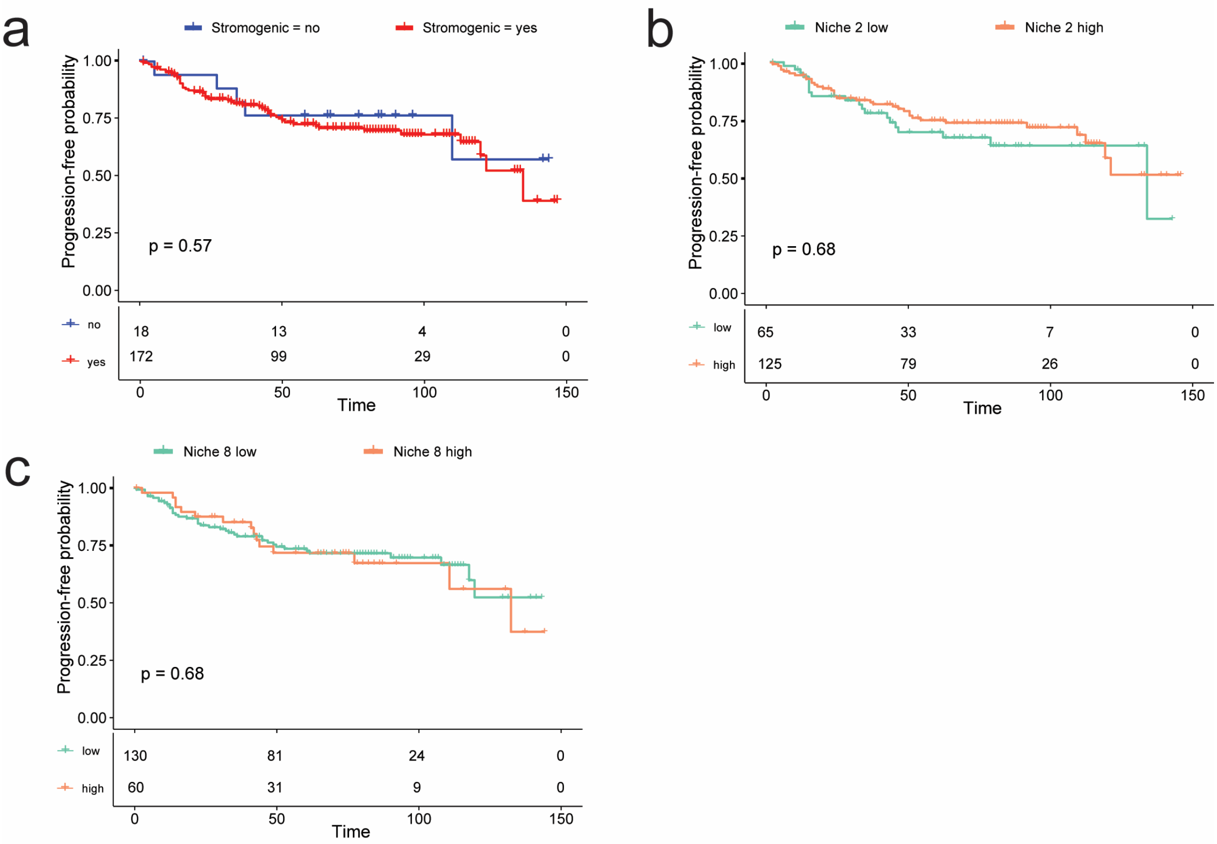


**Supplementary Figure 6. Niche 9 associates with worse clinical outcomes. a** Kaplan-Meier progression-free survival analysis with patients stratified by stromogenic status, defined by the presence of at least one core annotated as stromogenic; *p*-values were computed using a two-sided log-rank test. **b-c** Kaplan-Meier progression-free survival analysis for abundance of niche 2 and 8, respectively. Patients were grouped into high and low categories based on niche abundance, defined at the core level using the cohort median and aggregated to the patient level (high if ≥1 high-abundance core); p-values were computed using a two-sided log-rank test.

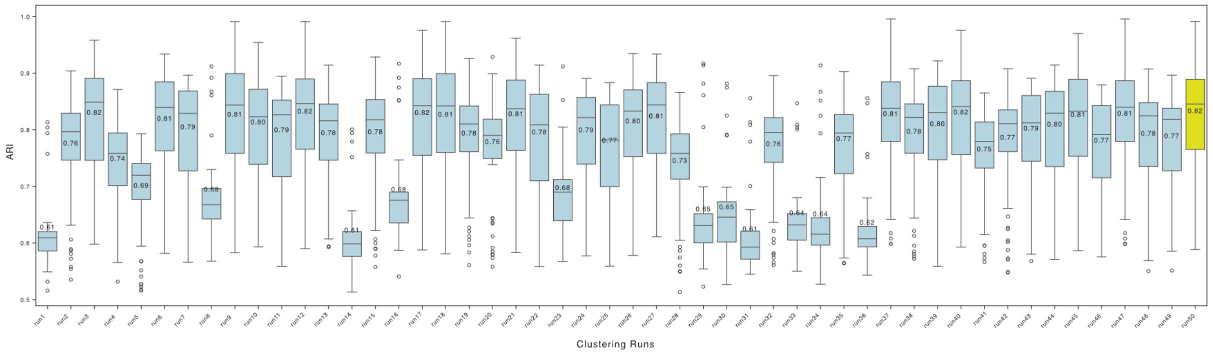


**Supplementary Figure 7. Robustness of tissue niche clustering assessed by adjusted Rand index (ARI).** K-means clustering was repeated 50 times using different random seeds. For each run, clustering agreement with all other runs was quantified using the adjusted Rand index (ARI). Each boxplot shows the distribution of pairwise ARI values between one clustering solution and all other runs. Higher ARI values indicate greater similarity between clustering solutions. The clustering run with the highest overall agreement across all comparisons is highlighted in yellow and was selected as the final solution for downstream analyses.

**Supplementary Table 1.** Description of the IMC antibody panel used.

| Protein target | Metal isotope | Company | Antibody clone | Product number | Dilution used |
|---|---|---|---|---|---|
| α-smooth muscle actin (α-SMA) | 141Pr | Standard BioTools | 1A4 | Maxpar Human Immuno-Oncology IMC Panlel Kit (#201508) | 1 to 400 |
| Prostate-specific antigen (PSA) | 142Nd | Cell Signaling | D11E1 | #36476F | 1 to 500 |
| Vimentin | 143Nd | Standard BioTools | D21H3 | Maxpar Human Immuno-Oncology IMC Panlel Kit (#201508) | 1 to 800 |
| Collagen type-I | 144 Nd | Standard BioTools | poly | 91H018144 | 1 to 200 |
| Synaptophysin (Syn) | 145Nd | Abcam | YE269 | ab187259 | 1 to 100 |
| Cytokeratin-5 (KRT5) | 146Nd | Abcam | EP1601Y | ab214586 | 1 to 800 |
| Phosphorytated-YAP1 (phospho Y357) | 147Sm | Abcam | EPR23680-48 | AB283327 | 1 to 25 |
| Pan-keratin | 148Nd | Standard BioTools | C11 | Maxpar Human Immuno-Oncology IMC Panlel Kit (#201508) | 1 to 200 |
| Carboxylesterase 1 (CES1) | 149Sm | Abcam | EP1376Y | AB235996 | 1 to 200 |
| Early growth response 1 (EGR1) | 150Nd | Abcam | EPR15916 | AB232448 | 1 to 100 |
| CD31 | 151Eu | Standard BioTools | EPR3094 | 3151025D | 1 to 200 |
| CD45 | 152 Sm | Standard BioTools | D9M81 | 3152018D | 1 to 400 |
| CD44 | 153 Sm | Standard BioTools | IM7 | 3153029D | 1 to 200 |
| FoxP3 | 155Gd | Standard BioTools | PCH101 | Maxpar Human Immuno-Oncology IMC Panlel Kit (#201508) | 1 to 200 |
| CD4 | 156Gd | Standard BioTools | ERP6855 | Maxpar Human Immuno-Oncology IMC Panlel Kit (#201508) | 1 to 400 |
| E-cadherin | 158Gd | Standard BioTools | 24E10 | Maxpar Human Immuno-Oncology IMC Panlel Kit (#201508) | 1 to 400 |
| CD68 | 159Tb | Standard BioTools | KP1 | Maxpar Human Immuno-Oncology IMC Panlel Kit (#201508) | 1 to 800 |
| CD66b | 160Gd | Standard BioTools | BLR111H | 91H033160 | 1 to 200 |
| CD20 | 161Dy | Standard BioTools | H1 | Maxpar Human Immuno-Oncology IMC Panlel Kit (#201508) | 1 to 800 |
| CD8a | 162Dy | Standard BioTools | C8/144B | Maxpar Human Immuno-Oncology IMC Panlel Kit (#201508) | 1 to 200 |
| CD11b | 163 Dy | Standard BioTools | EPR1344 | 91H007163 | 1 to 400 |
| Tumor Protein p63 (p63) | 164Dy | Abcam | EPR5701 | ab214790 | 1 to 50 |
| β-Catenin | 165Ho | Standard BioTools | D10A8 | 3165040D | 1 to 200 |
| Podoplanin (PDPN) | 166Er | Biolegend | NC-08 | 337002 | 1 to 200 |
| Endoglin (CD105) | 167Er | Abcam | EPR10145-12 | AB271922 | 1 to 400 |
| Ki-67 | 168Er | Standard BioTools | B56 | Maxpar Human Immuno-Oncology IMC Panlel Kit (#201508) | 1 to 200 |
| Tumor Protein p53 (p53) | 169Tm | Abcam | DO1 | ab237976 | 1 to 200 |
| CD3 | 170Er | Standard BioTools | Polyclonal | Maxpar Human Immuno-Oncology IMC Panlel Kit (#201508) | 1 to 200 |
| ETS Transcription Factor ERG | 171Yb | Abcam | EPR3864(2) | ab174739 | 1 to 50 |
| cleaved Caspase 3 (cCasp3) | 172Yb | Standard BioTools | 5A1E | 3172027D | 1 to 200 |
| Calponin 1 (CNN1) | 173Yb | Abcam | EP798Y | AB216651 | 1 to 6400 |
| Cytokeratin-8/18 (KRT8-18) | 174Yb | Cell Signaling | C51 | #51274F | 1 to 400 |
| Melanoma cell adhesion molecule (MCAM or CD146) | 175Lu | Abcam | EPR3208 | AB210072 | 1 to 100 |
| Androgen Receptor (AR) | 176Yb | Cell Signaling | D6F11 | #23791F | 1 to 200 |
| Nucleic acid (iridium) | 191Ir/193Ir | Standard BioTools | | Maxpar Human Immuno-Oncology IMC Panlel Kit (#201508) | 1 to 400 |
| Cell segmentation kit (ICSK 1/2/3) | 195Pt/196Pt/198Pt | Standard BioTools | | 201500 | 1 to 100 |
